# A Tissue-Targeted Immunotherapy Protects Against Recurrent Ocular Herpes Infection and Disease, and Alleviates the Associated Cognitive, Behavioral, and Social Deficits in a ‘Humanized’ HLA Transgenic Mouse Model of UVB-Induced Reactivation

**DOI:** 10.64898/2026.09.27.754856

**Authors:** Sweta Karan, Swayam Prakash, Aziz A. Chentoufi, Khan Intharachalit, Sarah Xue Le Ng, Junseong Hwang, Chhaya Maurya, Lbachir BenMohamed

## Abstract

**Background:** Recurrent ocular herpetic disease is a leading infectious cause of corneal blindness and is associated with psychological distress, anxiety, and social withdrawal. No ocular herpes vaccine or immunotherapy currently exists.

**Hypothesis:** A prime/pull/keep (PPK) T-cell immunotherapy, built on HSV-1 epitopes preferentially recognized by CD8^+^/CD4^+^ T cells from naturally protected asymptomatic (ASYMP) individuals, would reduce viral reactivation, shedding, recurrent corneal disease, and associated neurobehavioral deficits.

**Methods:** Immunodominant HLA-restricted CD8+/CD4+ epitopes from HSV-1 gD/gB, tegument proteins VP11/12/VP13/14, and helicase UL9 were delivered via topical ocular AAV8 vector (prime) to latently infected HLA-A*0201/DRB1 mice, combined with chemokine CXCL11 (pull) and IL-2/IL-15 cytokines to sustain TRM cells (keep), then UVB-exposed to trigger reactivation and recurrent corneal disease.

**Results:** PPK immunotherapy conferred robust, near-complete protection against recurrent HSV-1 disease and improved survival after UVB-induced reactivation, unlike its individual components. This protection was associated with site-specific accumulation of HSV-1-specific CD8^+^/CD4^+^ T^RM^ cells in the cornea and trigeminal ganglia (TG), but not in the spleen, with peak retention from combined IL-2/IL-15, alongside durable central/effector memory populations. PPK-induced T cells showed enhanced polyfunctionality (cytokine production, cytotoxicity, proliferation), reduced exhaustion markers, and a TCF-1+ self-renewing phenotype. PPK also elicited robust, Th1-biased HSV-1-specific IgG responses complementing mucosal immunity, and restored spatial memory, reduced depression-like behavior, and reversed social withdrawal in mice with recurrent ocular herpes.

**Conclusion:** PPK immunotherapy induces potent, durable antiviral T_RM_ cells at primary sites of HSV-1 latency and reactivation, protecting against recurrent ocular herpetic disease and alleviating associated neurobehavioral deficits, supporting its translational potential in humans.

**IMPORTANCE:** Recurring ocular herpes caused by HSV-1 is a major cause of blindness worldwide, and living with repeated flare-ups also takes a serious emotional toll, contributing to anxiety, low mood, and social withdrawal. Despite how common and damaging this condition is, no vaccine or treatment currently exists to prevent these outbreaks. This study tested a new three-part immune therapy designed to boost the body’s defenses against the virus, using viral pieces the immune system recognizes especially well in people who carry it without ever getting sick. In mice carrying human-like immune genes, this therapy trained immune cells to gather and stay active at the exact sites where the virus hides and can reawaken: the eye and nearby nerve tissue. Treated mice showed better protection from repeat outbreaks and improved survival after UV light triggered reactivation, compared to mice given only partial versions of the therapy. Treated mice also produced strong antibody responses and showed noticeable improvements in memory, mood, and social behavior after infection, suggesting the therapy protected the brain and behavior as well as the eye. These findings suggest this new therapy could offer meaningful protection against both the physical and emotional burden of recurrent ocular herpes in the clinic.

## INTRODUCTION

Herpes simplex virus type 1 (HSV-1) is a leading cause of infectious corneal blindness, affecting approximately 3.72 billion individuals globally (1). The mechanisms by which HSV-1 enters human corneal epithelial cells remain poorly understood, though they involve a complex interplay between viral glycoproteins and host cell receptors (2, 3). Corneal infection can occur during both primary and recurrent episodes. Following the initial infection, HSV-1 evades the immune system and establishes lifelong latency in sensory neurons, primarily within the trigeminal ganglion (TG) via the ophthalmic branch (V1) of the trigeminal nerve (4, 5). This latent viral reservoir can signal to orofacial mucosal tissues and is associated with an increased risk of viral reactivation, inflammatory responses, and overall clinical disease burden (6, 7). Potential corneal pathologies caused by HSV-1 include both asymptomatic and symptomatic herpes simplex keratitis, blepharitis, conjunctivitis, neovascularization, and disciform stromal edema (8). These conditions can progress to severe complications such as reduced corneal sensation, scarring, stromal opacity, and, ultimately, blindness (9). Herpes infection is also associated with significant psychological distress, including emotional burden, stigma, anxiety, and reduced quality of life (10). In many patients, the diagnosis and recurrent outbreaks may contribute to feelings of shame, social withdrawal, and emotional harm, and distress may in turn worsen recurrence risk (10). Due to the increasing prevalence and recurrence of HSV-1-associated corneal disease, the development of effective therapeutic strategies is critically important.

Current antiviral therapies, including corticosteroids and nucleoside analogs, aim to suppress ocular inflammation and inhibit viral replication (11). However, these treatments do not eliminate the latent virus; they primarily reduce viral shedding and relieve only about 45 % of herpetic disease symptoms, allowing corneal complications to persist (8, 12). Because of the limited efficacy of existing therapies and the significant public health burden of HSV-1 corneal infections, the development of an effective therapeutic vaccine represents a promising strategy for long-term disease control and prevention.

Several vaccine approaches, such as inactivated or attenuated vaccines, replication-defective vaccines, subunit-based protein, DNA, mRNA vaccines, and adeno-associated virus-based vaccines, have been investigated in animal models (mouse, guinea pig, cotton rat, and rhesus macaque) and human clinical studies. None have received licensure for use against herpes simplex virus (13, 14). This failure is due to an incomplete understanding of the immune responses required for adequate protection against HSV (15). Vaccines targeting HSV-specific antigens, including glycoproteins gB, gD, gC, and non-glycoproteins, have been shown to induce high levels of neutralizing antibodies and stimulate adaptive cellular immune responses (14, 16). However, these strategies have ultimately failed to provide sufficient protection in both prophylactic and therapeutic settings. Multi-epitope T cell-based vaccines provide improved therapeutic and protective efficacy by eliciting robust memory T cell responses (17–19).

T cells play a critical role in the adaptive immune system by controlling infections, maintaining immune surveillance, and influencing the outcomes of vaccines and immunotherapies (20, 21). Upon antigen stimulation, naïve T cells differentiate into a diverse population of effector and memory cells, characterized by their phenotype, functional capacity, and differentiation potential (22). These include short-lived effector CD8⁺ T cells (T_EFF_), exhausted CD8⁺ T cells (Tex), long-lived memory CD8⁺ T cells (T_LM_), memory precursor CD8⁺ T cells (T_MP_), central, effector, and tissue-resident memory T cells (T_CM_, T_EM_, T_RM_) (23). T_RM_ cells have emerged as key players in mediating site-specific, long-lasting immunity at peripheral tissues (24, 25). Emerging evidence suggests that stimulation of mucosal immune responses induces tissue-resident memory cells (T_RM_), providing superior and localized protection compared to circulating memory T cells (26, 27). T_RM_ cells exist within non-lymphoid tissues, including the trigeminal ganglion, and promote targeted responses upon re-infection(28). The generation and maintenance of T_RM_ cells is strongly regulated by chemokines and cytokines that govern T cell recruitment, localization, and differentiation (29–31). The impact of intestinal epithelial cells-derived CXCL11 has been reported to drive T cell migration towards IFN-γ-treated intestinal epithelium, enhancing anticancer T cells. It is reported to improve anticancer T cells within the tumor microenvironment (32). Similarly, IL-2 influences terminal effector CD8 T cell differentiation, while IL5 is reported to promote memory cell phenotype. Both IL-15 and IL-2 play distinct but complementary roles in maintaining tissue-resident memory (T_RM_) cells (33).

Adeno-associated viral vectors are currently among the most effective and safe tools for gene delivery, including chemokines and cytokines, due to their broad tissue tropism and promising safety profile (34). The AAV8 vector can stably transduce tissues such as the brain, glial cells, and neuronal tissues, resulting in 10-to-100-fold higher expression of the target gene compared to other serotypes (35, 36). Several AAV8-mediated transgenes, such as LDLR, G6PC, FVIII, and RLBP1, are currently in clinical trials (35, 37–39).

Both the trigeminal ganglia (TG) and the cornea play central roles in HSV-1 pathogenesis, serving as primary sites for latency, reactivation, and acute infection, respectively (40). The limited expression of T cell-attracting chemokines such as CXCL9, CXCL10, and CXCL11 in non-infected TG impairs the recruitment of antiviral CD8⁺ T cells from the circulation (41, 42). HSV-1-specific CD8⁺ T cell epitopes assist the migration and retention of T_RM_ cells at the critical sites of HSV-1 infection, including cornea and TG, enhancing immune surveillance (43, 44). Therefore, to enhance T_RM_ response at mucosal sites, ‘prime/pull’ and ‘prime/pull/keep’ strategies have been developed. The present study focused on the ‘prime/pull/keep’ T cell immunotherapy approach which involves an initial systemic priming with HSV-1 antigen encoding for ten asymptomatic (ASYMP) CD8^+^ T-cell peptide epitopes and 3 CD4^+^ T-cell epitopes from the glycoproteins D and B (gD and gB), viral tegument proteins (VP11/12 and VP13/14) and the DNA replication binding helicase (UL9), all specially recognized by CD8^+^ and CD4^+^ T cells from “naturally protected” HSV-1-seropositive healthy ASYMP individuals antigens to activate T cells, followed by the localized application of chemokines-CXCL11 to recruit these cells to the target tissue (pull). Subsequently, cytokines such as IL-2 and IL-15 are applied to promote the maintenance of T_RM_ cells within the tissue (keep). This strategy thereby establishes a robust T cell immunity within the trigeminal ganglion, offering a long-term protective response against HSV-1 ocular infection.

## MATERIALS AND METHODS

### AAV8 viral prime/pull/keep (PPK) T cell immunotherapy construct

To induce robust T cell responses, asymptomatic (ASYMP) CD8⁺ and CD4⁺ T cell peptide epitopes were screened and identified from HSV-1 glycoproteins D and B (gD and gB), viral tegument proteins (VP11/12 and VP13/14), and the DNA-binding helicase UL9, as previously described (45, 46). All peptides were synthesized by 21st Century Biochemicals (Marlboro, MA) and purified to 95–98% purity using reversed-phase high-performance liquid chromatography (Vydac C18), with confirmation by mass spectrometry (Voyager MALDI-TOF System). Stock solutions were prepared at a concentration of 1 mg/mL in PBS, aliquoted, and stored at –20 °C. The prime/pull/keep (PPK) AAV8 viral vectors expressing 10 CD8⁺ and 3 CD4⁺ T cell epitopes, the chemokine CXCL11, and cytokines IL-2 and/or IL-15 under the control of the neurotropic Ca²⁺/calmodulin-dependent protein kinase II subunit alpha (CamKIIα) promoter were synthesized by Vector Biolabs, Inc., as described earlier (46). Distinct CD8⁺ T cell epitopes were separated using the AAY linker, while CD4⁺ T cell epitopes were separated by the GPGPG linker, as shown in Figure 1. The prime AAV8 vector encodes only the selected T cell epitopes. The prime/pull-AAV8 vector co-expresses the T cell epitopes and mouse CXCL11 chemokine. The prime/pull/keep AAV8 vector expresses the T cell epitopes, mouse-specific CXCL11 chemokine, and either mouse IL-2, mouse IL-15, or a combination of both cytokines.

**Figure 1.**
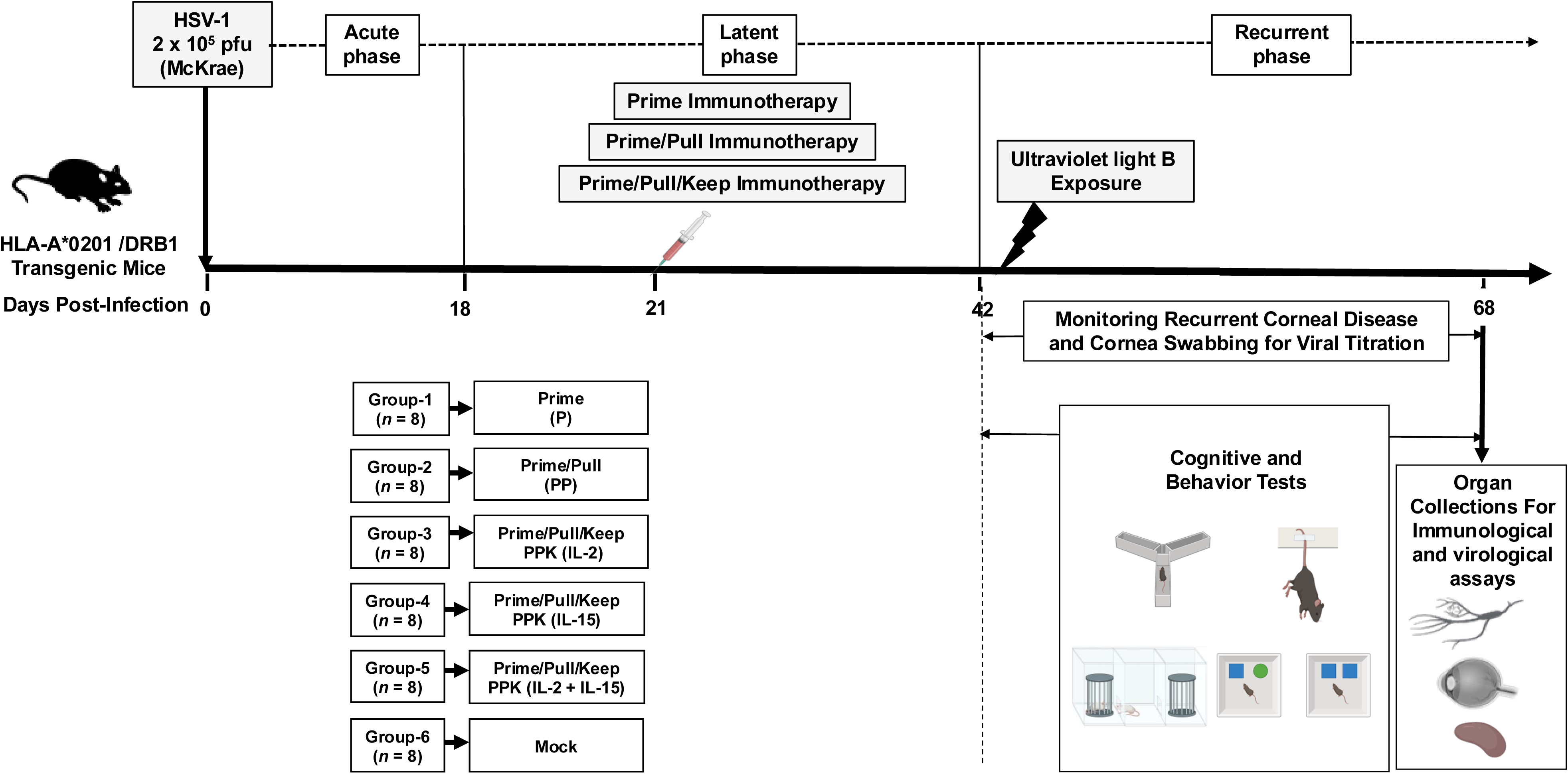
Experimental outline of the therapeutic prime/pull/keep T cell immunotherapy in HSV-1-infected HLA-A*0201/DR-B1 transgenic mice model: Double-transgenic mouse model of ocular herpes (HLA-A*0201/DRB1) Tg mice HLA transgenic mice were subjected to infection with HSV-1 (strain McKrae; 2 × 10⁵ PFU/eye) on day 0 and monitored for an acute phase of infection lasting up to 18 days. Post-acute-infection, mice were treated with adeno-associated viral vectors (AAV8) based PPK T cell immunotherapy (1 × 10⁷ genome copies). Upon UVB treatment, corneal disease and viral reactivation were monitored during the latent phase of herpes infection between 42-62 days. At the endpoint, the mice were euthanized, and trigeminal ganglia, corneas and spleen were collected from the mock, prime, prime/pull, and prime/pull/keep immunotherapy treated groups. The selected CD8⁺ and CD4⁺ T cell epitopes from the HSV-1 major antigens gD, gB, VP11/12, VP13/14, UL9 and linker sequences are indicated in bold. The prime AAV8 vector encodes only the selected T cell epitopes. The prime/pull AAV8 vector co-expresses the epitopes along with mouse CXCL11 chemokine. The prime/pull/keep AAV8 vector expresses the epitopes, CXCL11, and either mouse IL-2, IL-15, or a combination of both cytokines. P represents the prime-only group receiving antigen priming without subsequent pull or keep phases, serving as a minimal immunization control. PP represents the prime/pull group expressing T cell-attracting chemokine CXCL11. PPK (IL-15) represents the prime/pull/keep immunotherapy group receiving IL-15 alone. PPK (IL-2) represents the prime/pull/keep immunotherapy group receiving IL-2 alone. PPK (IL-2 + IL-15) represents the prime/pull/keep immunotherapy group receiving combined IL-2 and IL-15 cytokines to enhance T cell survival, proliferation, and tissue retention. NI represents the non-immunized control group with no treatment or antigen exposure. PI represents pre-immune sera collected before infection.

### T cell-based immunotherapy in HLA-A*0201/DR-B1 transgenic mice

Prime/Pull/Keep immunotherapy was performed in HLA-A*0201/DR-B1 transgenic mice after receiving approval from the Institutional Animal Care and Use Committee at the University of California, Irvine (UCI), and followed the Guide for the Care and Use of Laboratory Animals published by the US National Institutes of Health (IACUC protocol # AUP-22-086). HLA-A*0201/DR-B1 transgenic mice, initially provided by Dr. François Lemonier (Pasteur Institute, Paris), were bred and maintained at the UCI. HLA-A*0201/DR-B1 molecule presents the immunogenic peptide epitopes distinguishing symptomatic and asymptomatic herpes responses. Age (8-week-old) and gender matched HLA-A*0201/DR-B1 transgenic mice (*n* = 48 total; 8 mice in each group) were ocularly infected with 2 × 10^^5^ PFU of HSV-1 (strain McKrae) on day 0. The viral suspension was prepared in 2.5 µl of DMEM and administered as an eye drop after corneal scarification. Post-acute infection, mice were treated by topical ocular administration of 1 x 10^7^ viral genome copies (GC) of a recombinant prime AAV8 vector alone (P), a prime/pull AAV8 vector (PP), or a prime/pull/keep AAV8 vector expressing murine IL-2 (PPK1: PPK IL-2), IL-15 (PPK2: PPK IL-15), or a combination of both cytokines (PPK3: PPK IL-2+IL-15). The control mock group was treated with the empty AAV8 vector.

### Herpes simplex virus production

HSV-1 (strain McKrae) was purified by ultracentrifugation through a sucrose gradient and titrated using the triple plaque assay as described earlier (47). The virus was passaged in rabbit skin cell monolayers in MEM (1×) supplemented with 10 % heat-inactivated FBS, 2 mM L-glutamine, 2.5 μg/ml amphotericin, and 5 % penicillin-streptomycin solution.

### Ultraviolet-B light-induced reactivation of latent HSV-1

Reactivation of latent HSV-1 infection was induced by UVB exposure 42 days post-infection, as described earlier (48). UVB light was emitted at a peak wavelength of 302 nm using a UVP Chromato-VUE TM20 transilluminator (UVP, San Gabriel, CA). Mice were anesthetized by intraperitoneal injection of ketamine/xylazine (0.1 mL per 20 g body weight, containing 87.5 mg/kg ketamine and 12.5 mg/kg xylazine). The eyes of the mice were exposed to UVB light at a dose of 250 mJ/cm² for 60 seconds on the transilluminator.

### Monitoring of recurrent ocular herpes infection and disease

Post-T-cell immunotherapy and UVB reactivation, the mice were monitored for ocular herpes virus infection and disease progression. Eyes were swabbed daily for up to 14 days using moist type 1 calcium alginate swabs, which were then frozen at -80 °C until titrated. The recurrent corneal herpetic disease was examined by fluorescein staining followed by imaging using a Leica MZ10F fluorescent stereomicroscope. Disease scoring in mice was recorded on a 0 - 4 scale (0 = no disease; 1 = 25%; 2 = 50%; 3 = 75%; 4 = 100%) as previously described (45).

### Quantification of ocular infectious virus

The HSV-1 viral titers were determined by real-time PCR using SYBR Green Master Mix (ThermoFisher A25742) and HSV-1 specific primers (Forward Primer: 5’-CATCACCGACCCGGAGAGGGAC-3’ and Reverse Primer: 5’-GGGCCAGGCGCTTGTTGGTGTA-3’) as described earlier (49). For absolute quantification, a standard curve was generated using the Amplitron® HSV-1 DNA Control (Vercelli, Granada, Spain). The lyophilized genomic DNA control was reconstituted in molecular biology-grade water according to the manufacturer’s instructions to achieve a stock concentration of 1.5×10^4^ copies/µl. Serial ten-fold dilutions of the reconstituted genomic DNA were prepared fresh in DNA-free water to establish a standard reference curve from 10^1^ to 10^5^ viral copies per reaction. Amplification of both the standard dilution series and experimental samples was performed in triplicate on an Applied Biosystems QuantStudio 5 quantitative real-time PCR, and absolute HSV-1 copy numbers were calculated by interpolating cycle threshold (CT) values against the standard curve.

### Isolation of corneal and trigeminal ganglion (TG) resident leukocytes and splenocytes in HSV reactivated and treated mice

At the study endpoint, mice from all experimental groups were euthanized, and the trigeminal ganglia (TG), cornea, and spleens were harvested individually. Corneal and TG tissues were digested in complete RPMI 1640 medium supplemented with 10% fetal bovine serum (FBS) and 2.5 mg/mL collagenase type IV (ThermoFisher Scientific17104019). Tissue digestion was performed at 37°C for 30 min with continuous shaking. Following digestion, the cell suspensions were passed through sterile 70 μm nylon cell strainers and washed with RPMI 1640 medium containing 10% FBS. Leukocytes were then enriched by density gradient centrifugation using 70% Percoll. Single-cell suspensions from spleens were prepared by mechanical dissociation through 70 μm cell strainers, followed by red blood cell lysis using ACK lysis buffer. Cells were washed, viability was determined by Trypan blue, and stained with fluorochrome-conjugated anti-mouse antibodies for flow cytometric analysis.

### Flow cytometry: Tetramer staining followed by surface and intracellular staining

Single-cell suspensions from the trigeminal ganglia (TG), cornea, and spleen were used for tetramer staining with HLA-A2-peptide tetramers provided by the NIH Tetramer Facility (50, 51). The peptide sequences corresponding to each HLA-A2 tetramer are listed in Supplementary Table 1. Briefly, cells were incubated with 1 μg/ml of each PE-labeled HLA-A2-peptide tetramer at 37°C for 30-45 min. Following tetramer staining, surface staining was performed using fluorochrome-conjugated anti-mouse antibodies specific for CD8 53-6.7, CD4 GK1.5, CD62L MEL-14, CD44 IM7, CD49a HMα1, CD103 S18001A, CD69 H1.2F3, CD107a 1D4B, PD-1 29F.1A12, TIM-3 B8.2C12, TIGIT 1G9, and LAG-3. C9B7W and CD39 GK1.5 (Biolegend or BD Pharmingen). Cells were then fixed and permeabilized using the appropriate fixation/permeabilization buffer according to the manufacturer’s instructions. Intracellular staining was subsequently performed in 1× Perm/Wash buffer on ice using antibodies against IFN-γ XMG1.2, Ki-67 16A8, Granzyme B QA18A28, TNF-α MP6-XT22, TCF-1 S33-966, TOX NAN448B, EOMES X4-83, and CTLA-4 63828. After staining, cells were washed and resuspended in 150 μl of BD Stabilizing Fixative buffer before acquisition. A minimum of 200,000 events per sample were acquired using a BD LSRFortessa flow cytometer and analyzed with FlowJo software, version 10.10 (FlowJo INC., Ashland, OR). For compensation controls, single-stained antibody capture beads (BD Biosciences) were used for each fluorophore. Fluorescence-minus-one (FMO) controls were included to establish gating boundaries, and known negative cell populations were used to verify background fluorescence and optimize gating strategies (**Supplementary Fig 1**). Briefly, lymphocytes were first identified based on forward and side scatter properties, followed by gating on singlets, viable cells (Zombie Aqua-negative), CD4^+^ and CD8^+^ T-cell populations, and finally tetramer-positive HSV-1-specific CD8^+^ T cells.

### Immunohistochemistry of cornea and trigeminal ganglia

Corneas and trigeminal ganglia were harvested on day 100, embedded in Tissue-Tek OCT compound (VWR International, West Chester, PA), and snap-frozen. Cryosections approximately 10 µm thick were prepared, fixed in acetone, and stored at –80°C until staining. Longitudinal sections were stained with either hematoxylin and eosin (H&E) or with antibodies specific for tissue-resident memory (T_RM_) cell markers-Alexa Fluor® 488 anti-mouse CD4 antibody (Biolegend-100423), PerCP anti-mouse CD8a antibody (Biolegend-100732), PerCP/Cyanine5.5 anti-mouse CD103 antibody (Biolegend-121416). To detect HSV-1 viral antigen, sections were incubated with HSV type 1 monoclonal antibody (Invitrogen-MA5-16285) followed by secondary staining with Goat anti-Mouse IgG (H+L), Alexa Fluor™ 488 antibody (Invitrogen-A28175). Imaging was performed using a Keyence BZ-X700 microscope at 20× magnification.

### Enzyme-linked immunosorbent assay (ELISA)

Serum samples were collected at the study endpoint from PPK T cell immunotherapy-treated and untreated HLA-A*0201/DRB1 transgenic mice and analyzed for HSV-1-specific antibody responses against HSV-1 glycoprotein D (gD) and whole HSV-1 virions by enzyme-linked immunosorbent assay (ELISA). Nunc 96-well flat-bottom maxisorp immune plates (ThermoFisher 442404) were coated overnight at 4°C with either 100 ng/well of recombinant HSV-1 glycoprotein D (Sino Biological, USA) or heat-inactivated HSV-1 virions (1 × 10⁵ PFU per well). Plates were subsequently washed and blocked with 3% bovine serum albumin (BSA) in PBS for 2 h at room temperature (RT). Following blocking, serial dilutions of mouse sera (1:20 to 1:312,500; 100 μl/well) were added and incubated overnight at 4°C. Plates were then washed three times with PBS containing 0.05% Tween-20 (PBST). Bound antibodies were detected using horseradish peroxidase (HRP)-conjugated secondary antibodies specific for mouse total IgG, IgG1, IgG2a, IgG2b, IgM, and IgA (all used at a 1:5,000 dilution; Invitrogen, USA) and incubated for 1h at RT. After washing, 100 μl/well of tetramethylbenzidine (TMB) substrate solution (Thermo Fisher Scientific, USA) was added, and the reaction was allowed to develop for 10–20 min in the dark. The reaction was terminated by the addition of 100 μl/well of stop solution (Thermo Fisher Scientific, USA), and absorbance was measured at 450 nm using a microplate reader. Antibody isotype profiling (IgG1, IgG2a) was conducted to characterize the Th1 versus Th2 polarization of the humoral immune response. The cutoff value for seropositivity was defined as the mean optical density (OD) of negative control sera plus three standard deviations (SD).

### Behavior studies

To assess the neurological and physiological impact of ocular reactivation in PPK T cell immunotherapy-treated and untreated mice, behavioral assays were performed as described earlier (52).

### The Y-maze spontaneous alternation test

Spatial working and reference memory were assessed using the Y-maze test (53, 54). As the recurrent HSV-1 infection has been associated with neuroinflammation and cognitive impairment, the Y-maze was used to evaluate the impact of HSV-1 reactivation and PPK immunotherapy on memory performance. For spontaneous alternation testing, mice were allowed to freely explore all three arms of the maze for 8 min. The sequence of arm entries was recorded, and the percentage of spontaneous alternation was calculated as an index of spatial working memory. To evaluate spatial reference memory, one arm of the maze was blocked during the training phase, allowing mice to explore the remaining two arms. After a 30-60 min retention interval, mice were returned to the maze with all arms accessible. The time spent and number of entries into the previously blocked (novel) arm were recorded as measures of spatial memory performance.

### Tail suspension test

Depression-like behavior associated with chronic HSV-1 infection and recurrent viral reactivation was evaluated using the tail suspension test (TST) (55). Mice were individually suspended by the tail using adhesive tape affixed approximately 1 cm from the tip of the tail and monitored for 6 min. The total duration of immobility was recorded and used as an indicator of behavioral despair and neuropsychological impairment. Increased immobility time was interpreted as enhanced depression-like behavior, whereas decreased immobility reflected improved behavioral performance and stress-coping responses.

### Novel object location (NOL) test

Spatial memory was assessed using the novel object location (NOL) test (52). Because recurrent HSV-1 infection can impair hippocampal-dependent cognitive function through chronic neuroinflammation, the NOL test was used to evaluate the effects of HSV-1 reactivation and PPK immunotherapy on spatial memory. Mice were first habituated to an open-field arena for 10 min. Twenty-four hours later, mice were allowed to explore two identical objects for 10 min (familiarization phase). After a 3 h retention interval, one object was relocated to a novel position, and mice were allowed to explore the arena for 10 min. Exploration time directed toward the relocated object was recorded, and a location discrimination index was calculated as an indicator of spatial memory performance.

### Novel object recognition (NOR) test

Recognition memory was evaluated using the novel object recognition (NOR) test (52). Following the NOL assessment, mice were re-exposed to the arena 24 h later, where one of the familiar objects was replaced with a novel object while maintaining the same object locations. Mice were allowed to explore the objects for 5 min, and the time spent investigating the novel and familiar objects was recorded. Recognition memory was quantified using a discrimination index calculated as the time spent exploring the novel object divided by the total exploration time of both objects. Increased preference for the novel object was interpreted as intact recognition memory, whereas reduced preference indicated HSV-1-associated cognitive impairment.

### Three-chamber sociability test

Social behavior and sociability were assessed using the three-chamber sociability test (56). Because recurrent HSV-1 infection can induce neuroinflammation and behavioral abnormalities, this assay was used to evaluate the effects of HSV-1 reactivation and PPK T cell immunotherapy on social interaction. During the test, mice were allowed to freely explore three interconnected chambers containing either a novel mouse or an empty enclosure. The time spent interacting with the novel mouse and the time spent in each chamber were recorded. Increased interaction with the social stimulus was interpreted as normal sociability and social motivation, whereas reduced social preference indicated HSV-1-associated neurobehavioral impairment.

### Three-chamber social novelty test

Social recognition memory was assessed using the three-chamber social novelty test (57). Because recurrent HSV-1 infection can affect cognitive and social behaviors through persistent neuroinflammation, this assay was used to evaluate the impact of HSV-1 reactivation and PPK immunotherapy on social memory. Following the sociability phase, mice were allowed to choose between a familiar mouse and a novel, previously unencountered mouse placed in separate chambers. The time spent interacting with each mouse was recorded. A preference for the novel mouse was interpreted as intact social recognition memory, whereas reduced preference indicated impaired social cognitive function associated with HSV-1-induced neurological dysfunction.

### Statistical analyses

GraphPad Prism 10 version 10.4.1 (La Jolla, CA) was used for determining the analysis of variance (ANOVA) and Student’s t-test to compare differences between mock, prime, prime/pull, prime/pull/keep T cell immunotherapy-treated group as described earlier(45). All flow cytometry data were acquired on a BD LSRFortessa and analyzed using FlowJo software, version 10.10, FlowJo INC., Ashland, OR.

### Data availability

The gene sequences of HSV-1 antigens, chemokines, and cytokines used in our prime/pull/keep T cell immunotherapy construct were extracted from NCBI with the following gene IDs and accession numbers. HSV-1 glycoprotein B (gB) (330109; M21629.1), glycoprotein D (gD) (L09243.1; AAA19629.1), viral tegument protein VP11/12 (UL46) (2703413; NC_001806.2), viral tegument protein VP13/14 (UL47) (2703415; NC_001806.2) and the DNA replication binding helicase protein (UL9) (2703434; NC_001806.2), CXCL11 (56066; NM_019494.1), IL2 (16183; NM_008366.3), IL15 (16168; NM_001254747.2).

## RESULTS

### 1. Prime/Pull/Keep Immunotherapy Targeting Asymptomatic Immunodominant CD4^+^ and CD8^+^ Epitopes Confers Robust Protection Against Recurrent HSV-1 Infection and Disease

To enhance the protective and immunotherapeutic efficacy of the prime/pull/keep T cell strategy, we designed a multi-epitope prime/pull/keep T cell immunotherapy construct comprising asymptomatic (ASYMP) CD4^+^ and CD8^+^ T cell epitopes from glycoproteins D and B (gD and gB), viral tegument proteins (VP11/12 and VP13/14), and the DNA replication-binding helicase (UL9). These ASYMP epitopes were identified from individuals who harbor the HSV-1 virus without experiencing recurrent symptoms, suggesting a protective and HSV-1-specific immune response (58). These epitopes were selected as they displayed higher binding affinities for soluble HLA-A*0201/DR-B1, ensuring optimal presentation and immunogenicity. These selected epitopes were cloned under the CamKIIa promoter in the adeno-associated virus type 8 vector (AAV8) for expression in the neurotropic tissues. The AAV8 vector construct expressing multiple T cell epitopes comprises 10 CD8^+^ and 3 CD4^+^ T cell epitopes, which were separated by linker AAY and GPGPG, respectively. An additional vector consisting of co-expressing T cell-attracting chemokine CXCL11 or T cell-retaining cytokines, IL-2/IL-15, was developed. The vector construct and details of the epitope used in the prime/pull/keep vaccination study have been illustrated in **Fig. 1**.

### 2. PPK Immunotherapy, but Not Its Individual Components, Achieves Near-Complete Disease Resolution and Survival After UVB-Induced HSV-1 Reactivation

To evaluate the therapeutic efficacy of the prime/pull/keep strategy against recurrent ocular herpes, the transgenic mice were ocularly infected with HSV-1 (McKrae, 2 × 10 ^^5^ pfu/eye). Post-acute infection, the latently infected mice were treated with either prime/pull/keep, prime/pull, or prime therapy. After UVB-induced viral reactivation, the mice were monitored for ocular herpes recurrence for up to 14 days. We systematically trace the impact of adding each component of the PPK immunotherapy compared to the mock-treated controls. As shown in **Fig. 2A**, the mice treated with PPK immunotherapy (PPK1, P=0.0164; PPK2, P=0.0113; PPK3, P=0.009) showed significantly reduced near-zero corneal disease scores, reducing primary stromal keratitis and blepharitis, compared to the control group. In contrast, prime alone provides a mild reduction in disease severity, but it does not reach statistical significance (P=0.2613), and adding the ‘Pull’ phase using CXCL-11 lowers the disease score further but remains statistically non-significant (P=0.1565) therapy group did not exhibit a significantly reduced disease score **Fig. 2A**.

**Figure 2.**
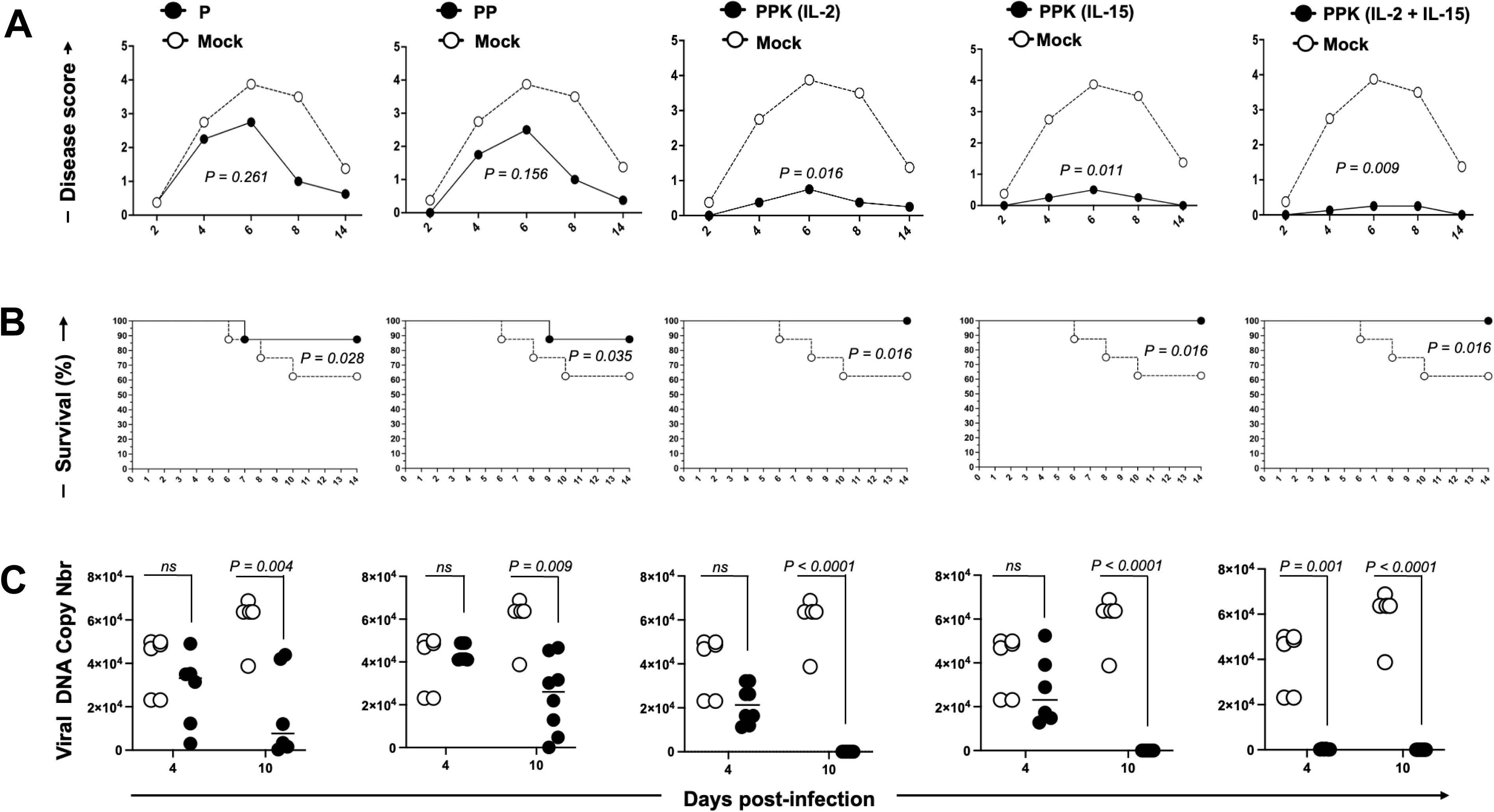
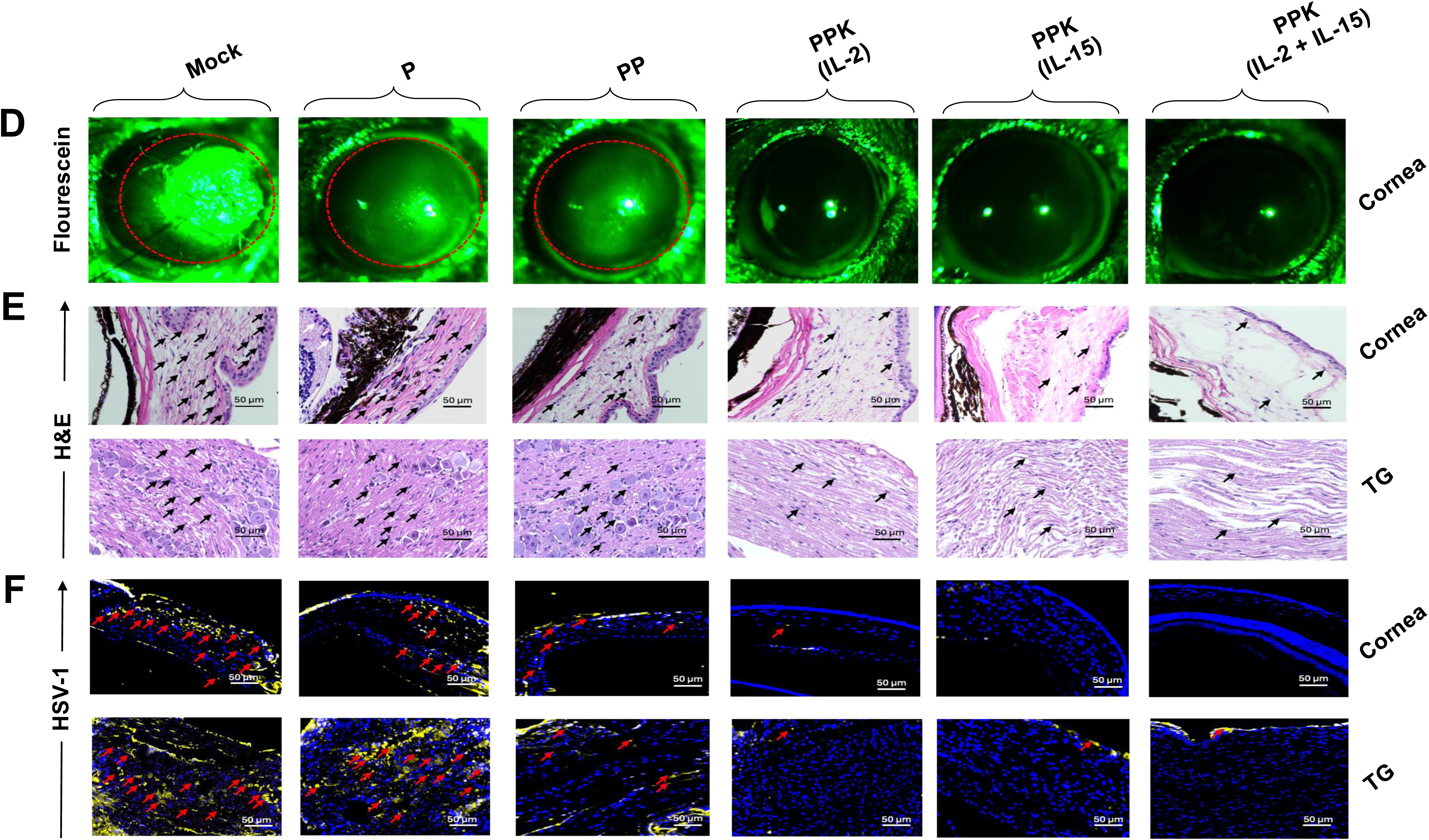
Prime/pull/keep T cell immunotherapy reduces recurrent corneal herpetic disease in HSV-1-infected HLA-A*0201/DRB1 Tg mice: **(A)** Recurrent corneal inflammation and cloudiness were assessed by scoring the severity of stromal keratitis and blepharitis. Stromal keratitis was scored as follows: 0, no disease; 1, mild cloudiness with visible iris details; 2, obscured iris details; 3, complete corneal opacity; and 4, corneal perforation. Blepharitis was scored as follows: 0, no disease; 1, puffy eyelids; 2, puffy eyelids with mild crusting; 3, eye swollen shut with severe crusting; and 4, complete eyelid closure due to swelling. Disease severity was significantly higher in the mock (untreated), prime only, and prime/pull groups compared with controls (*P < 0.05, unpaired t-test). (**B**) Mice were monitored for survival over a 14-day period following UVB treatment. Kaplan–Meier survival analysis showed 100% survival in the PPK immunotherapy group compared with 60% survival in the mock (untreated) group during the recurrent infection period (log-rank test, *P = 0.0168). Both the prime-only group (*P = 0.0281) and the prime/pull group (*P = 0.0356) exhibited 90% survival.

To evaluate the safety, efficacy, and immunogenicity of PPK T-cell therapy before human clinical trials, it is essential to assess the immune responses across various animal models. The ultraviolet B (UVB) induced herpes recurrence in HLA-A*0201/DR1 mice closely mimics environmental triggers mediated reactivation of HSV-1 in humans (59). The HLA-A*0201/DR1 mice express human HLA-A*0201 and HLA-DR molecules, thereby permitting antigen presentation and T cell activation within a humanized immunological framework (60–62). These mice develop recurrent herpetic stromal keratitis, blepharitis, and viral shedding, paralleling the clinical symptoms of human ocular herpes (8). We assessed the survival of PPK immunotherapy treated mice and observed robust 100% survival in all prime/pull/keep-treated groups (PPK1, PPK2, PPK3; P = 0.0168) in the presence of ‘Keep’ homeostatic cytokines either IL-2, IL-15, or the combined IL-2 plus IL-15 cocktails, compared to 90 % survival in the prime and prime/pull treated groups while the mock controls consistently drop down to approximately 60% survival following viral reactivation (**Fig. 2B**).

Then, we performed a comprehensive pathological evaluation of the ocular surface and underlying tissues following UVB reactivation. We monitored corneal lesions by fluorescein ocular surface staining. Fluorescein staining is a direct measure of corneal epithelial integrity, where bright green staining indicates active corneal ulceration and severe herpetic keratitis. We observed a progressive, though incomplete, reduction in the size of these lesions in the prime and prime/pull treated group.

PPK T-cell immunotherapy in the presence of ‘keep’ homeostatic cytokines -treated groups showed no green ulceration and complete preservation of the corneal epithelial sheet, indicating a significant reduction in corneal pathology, as indicated in representative **Fig. 2D**.

Then, we assessed HSV-1-associated pathology of the trigeminal ganglia and cornea by Hematoxylin and Eosin (H&E) staining. Hematoxylin and eosin (H&E) staining of trigeminal ganglia and corneal sections revealed reduced pathological damage, lesser inflammatory cell infiltration, preserved corneal stromal layers, and neuronal morphology in the trigeminal ganglia in the PPK immunotherapy-treated groups, indicating sustained protection against HSV-1-induced ocular disease. Mock control, prime, and prime/pull groups showed increased mononuclear cell infiltration and localized tissue disorganization. A dense influx of inflammatory cells into both the corneal stroma and the trigeminal ganglia mediates uncontrolled pathology, causing permanent tissue scarring and blindness (**Fig. 2E)**. PPK immunotherapy clears viral antigen and lowers HSV-1 genome copy number: We next wanted to confirm that this structural protection was directly associated with viral control. We assessed viral persistence in the tissue by direct immunofluorescence staining using HSV-1-specific antibodies. By utilizing IL-2, IL-15, or the combined IL-2 and IL-15 cytokines, PPK immunotherapy-treated groups showed reduced HSV-1 antigen presence in both the trigeminal ganglia and cornea, indicating clearance of latent recurrent virus. Mock control, prime alone, and prime/pull cohorts showed intense viral staining, indicating retention of reactivated viral replication throughout the tissue (**Fig. 2F**). Additionally, HSV-1 genome load and viral replication were quantified from the collected corneal swabs by real-time PCR using HSV-1-specific primers. Although all vaccinated groups containing homeostatic ‘keep’ cytokines maintained significantly lower viral DNA copy numbers, reduction in viral load and replication by day 10 (P < 0.005). Additionally, the PPK3 group co-administered with IL-2 plus IL-15 demonstrated a substantial reduction as early as day 4 (P = 0.001) (**Fig. 2C**). These results collectively demonstrate that the combination of the chemokine CXCL11 and cytokines IL-2/IL-15 prominently enhances the protective efficacy of the PPK T cell-based immunotherapy strategy by showing higher viral clearance, reducing HSV-1 replication, and reducing viral DNA copy numbers in latently infected tissues.

### 3. PPK Immunotherapy Drives Site-Specific Accumulation of HSV-1-Specific Tissue-resident memory CD4^+^ and CD8^+^ T_RM_ cells in the Cornea and Trigeminal Ganglia, with Peak Retention from Combined IL-2/IL-15

Next, we evaluated the underlying cellular immune response to therapeutic prime/pull/keep T cell immunotherapy. To determine whether PPK immunotherapy enhanced antigen-specific cellular immunity, we used MHC class I tetramer staining to assess the frequency and distribution of HSV-1-specific CD8^+^ T cells in the cornea, TG, and spleen using tetramers corresponding to the HSV-1 epitopes gD, gB, VP11/12, VP13/14, and UL9. Flow cytometric analysis revealed significantly higher frequencies of tetramer-positive CD8^+^ T cells in the homeostatic ‘keep’ cytokines-containing PPK-treated mice compared with untreated controls at the primary sites of HSV-1 reactivation and persistence, indicating efficient recruitment and maintenance of HSV-1-specific tissue-resident memory CD4^+^ and CD8^+^ T_RM_ cells at sites of latent infection. (**Figs. 4A** and **B**) This increase was greatest in the PPK group containing both cytokines, in both the cornea and TG. In this group, gB-specific tetramer-positive CD8⁺ T cells reached frequencies of ∼70% in the TG and ∼20.5% in the cornea, whereas only negligible background staining was detected in mock-immunized and negative control animals. The PPK groups receiving either IL-2 or IL-15 alone exhibited an intermediate level of T cell retention. The PP group demonstrated recruitment of cells to the tissue but lacked the enhanced long-term maintenance provided by the “Keep” (cytokine) signal. The prime group showed systemic priming, resulting in the lowest T cell frequencies among all the treated groups. At the same time, PPK- or PP-treated mice showed much lower and non-significant frequencies of HSV-1-specific CD8^+^ T cells in the spleen between ∼2% and ∼6%, suggesting the expansion and retention of CD8^+^ T cells at the targeted sites of infection, such as the corneas and the TG, rather than in the systemic compartment of the spleen (**Supplementary Fig 2B**).

### 4. PPK Immunotherapy Drives Site-Specific Accumulation of HSV-1-Specific Tissue-Resident Memory (T_RM_) Cells in the Cornea and Trigeminal Ganglia, but Not the Spleen

Then, we confirmed the expression of tissue-resident memory by staining classical T_RM_ cell surface markers: CD49a, CD69, and CD103 in the cornea, TG, and spleen. As we observed, Prime/Pull/Keep T cell immunotherapy enhances the retention and longevity of HSV-1-specific CD8^+^ T cells; these recruited T cells also showed increased expression of tissue-resident memory (T_RM_) T cell markers at sites of latency and infection. Tetramer-positive CD8⁺ T cells from PPK-treated mice exhibited significantly increased frequencies of HSV-1-specific T_RM_ cell markers, including CD69 (∼11% in TG and ∼20% in the cornea), CD103 (∼12% in both TG and cornea), and CD49a (∼14% in TG and ∼38% in the cornea), compared with untreated mock controls, prime alone, and prime/pull treated groups (**Figs. 3C** and **D**). In contrast, no significant increases in T_RM_ marker expression were observed in the spleen, where frequencies remained low (∼1.2–2%), indicating that PPK immunotherapy preferentially promotes the establishment and maintenance of tissue-resident antiviral immunity at sites of HSV-1 latency and reactivation (**Supplementary Fig. 2C**).

**Figure 3.**
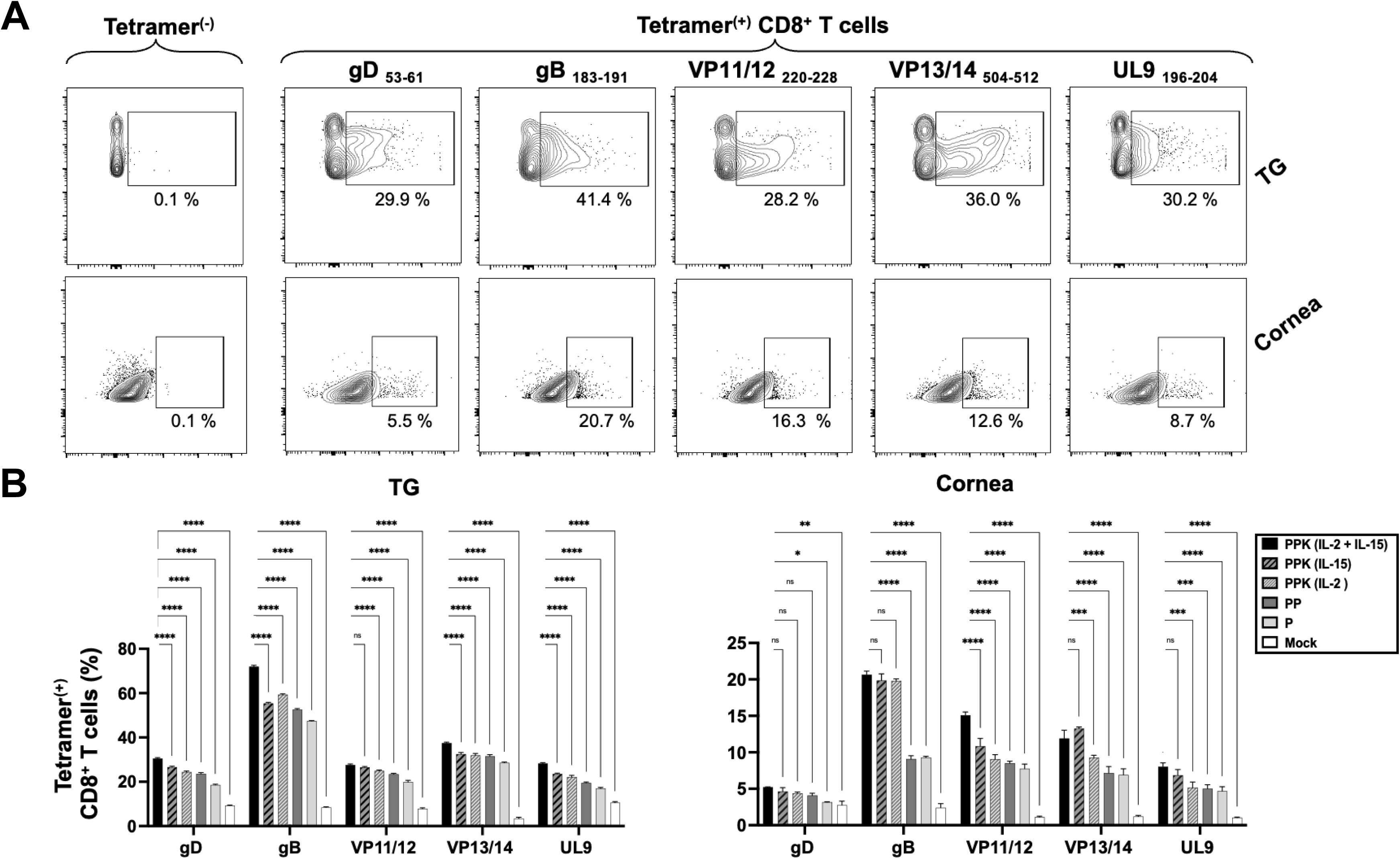
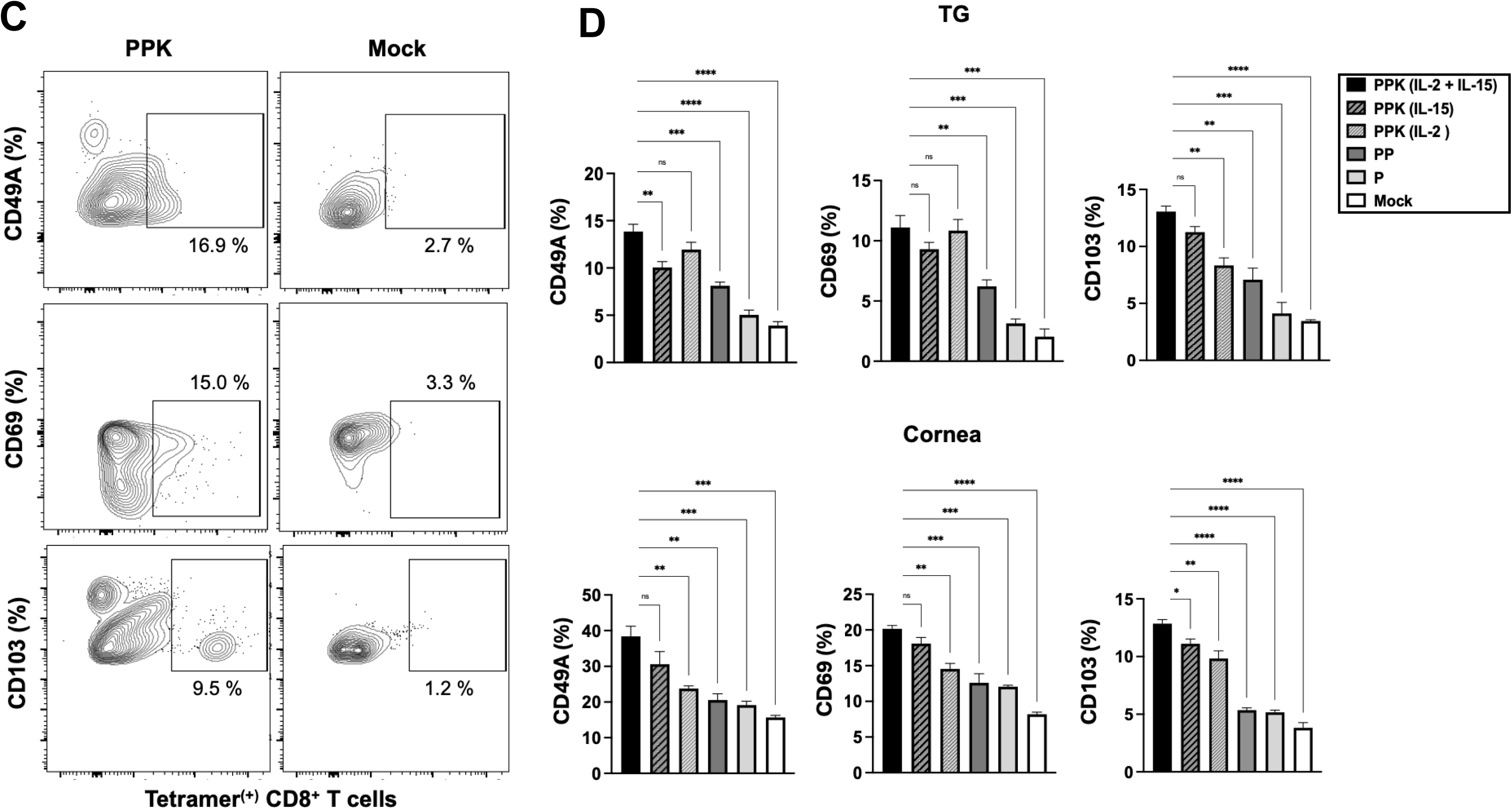
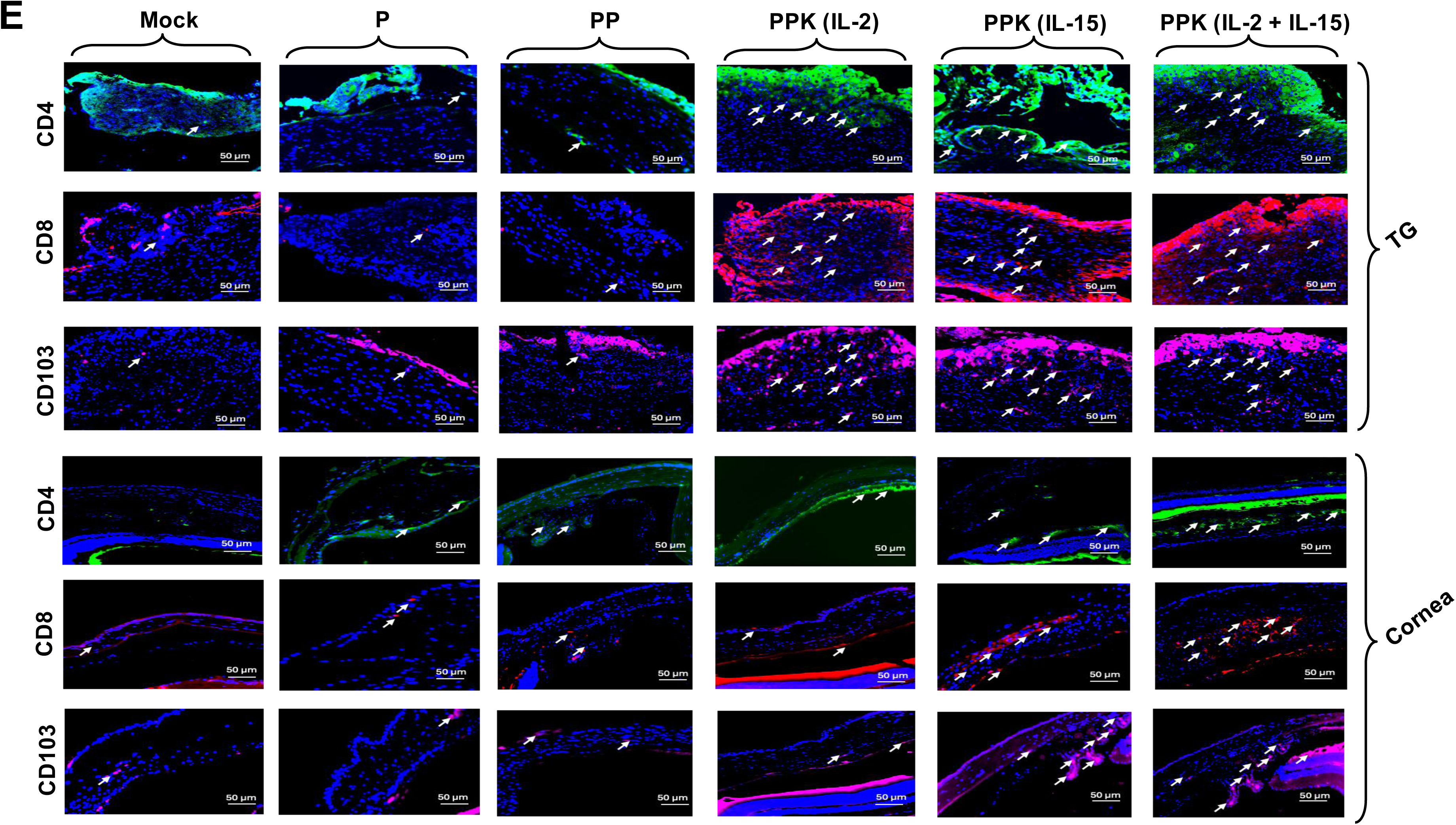
Prime/pull/keep T cell immunotherapy reduces pathology in HSV-1 infectedHLA-A*0201/DR-B1 transgenic mice: (**A**) Representative images showing reduced recurrent corneal herpetic disease in HLA-A*0201/DRB1 Tg mice treated with PPK immunotherapy in comparison to mock untreated HLA-A*0201/DRB1 Tg mice. The corneal lesions were visualized by fluorescein staining and imaged under a Leica MZ10F fluorescent stereomicroscope. (B) Representative images of Hematoxylin & Eosin (H&E) stained sections of the trigeminal ganglia and cornea from PPK T cell immunotherapy and mock untreated mice. Images were acquired at 40× magnification.

Then, we confirmed the tissue localization and infiltration of T_RM_ cells within the trigeminal ganglion and cornea by immunofluorescence microscopy. Immunobiological analysis revealed a significant increase in infiltrating CD4⁺, CD8⁺, and CD103⁺ T_RM_ cells in the PPK T cell immunotherapy-treated groups receiving the ‘Keep’ cytokines IL-2, IL-15, or their combination compared to the control, prime-only, and prime/pull treated groups (**Fig. 3E**). These findings correlate with our flow cytometric data, as shown earlier. The enhanced presence of T_RM_ cells supports the improved effectiveness of combining a T cell epitope-based prime therapy with the chemokine CXCL11 and cytokines IL-2 and IL-15, which likely promotes T_RM_ cell recruitment, retention, and long-term protective immunity. Taken together, these results demonstrate that the prime/pull/keep T cell immunotherapy incorporating specific T cell epitopes, T cell-attracting chemokine CXCL11, and T cell-retaining cytokines IL-2/IL-15 effectively induced CD8^+^ and CD4^+^ T cells that infiltrate T_RM_ cells in the trigeminal ganglia and cornea. These immune responses facilitated the clearance of HSV-1 virus and reduced HSV-1-associated pathology in both the trigeminal ganglia and cornea in the UVB-induced recurrent HSV-1 infection model.

### 5. PPK Immunotherapy Establishes Durable Central and Effector Memory CD8^+^ T Cell Populations Supporting Long-Term Protection Against HSV-1 Reactivation

Then, we evaluated the memory phenotype of these localized T cell populations in the trigeminal ganglion and cornea. We observed the differentiation of HSV-1-specific CD8^+^ T cells into central memory (T_CM_; CD44^+^CD62L^+^) and effector memory (T_EM_; CD44^+^CD62L^−^) subsets. Flow cytometric analysis demonstrated that PPK combining IL-2 and IL-15-treated mice exhibited significantly higher frequencies of both T_CM_ (∼22 % in TG, ∼35 % in cornea) and T_EM_ (∼22 % in TG, ∼25 % in cornea) populations compared with untreated controls, prime only, and prime/pull treated (**Figs. 4A** and **4B)**. The expansion of the T_CM_ population suggests the establishment of a durable memory pool capable of rapid proliferation upon antigen re-encounter, whereas the increased T_EM_ population indicates enhanced effector function for immediate antiviral responses. The elevated frequencies of memory CD8^+^ T cells were observed in the peripheral tissues, consistent with the ability of PPK immunotherapy to generate robust tissue-associated immune memory. Thus, these findings demonstrate that PPK immunotherapy promotes the development of long-lasting HSV-1-specific memory responses for sustained immune surveillance and protection against recurrent HSV reactivation.

**Figure 4.**
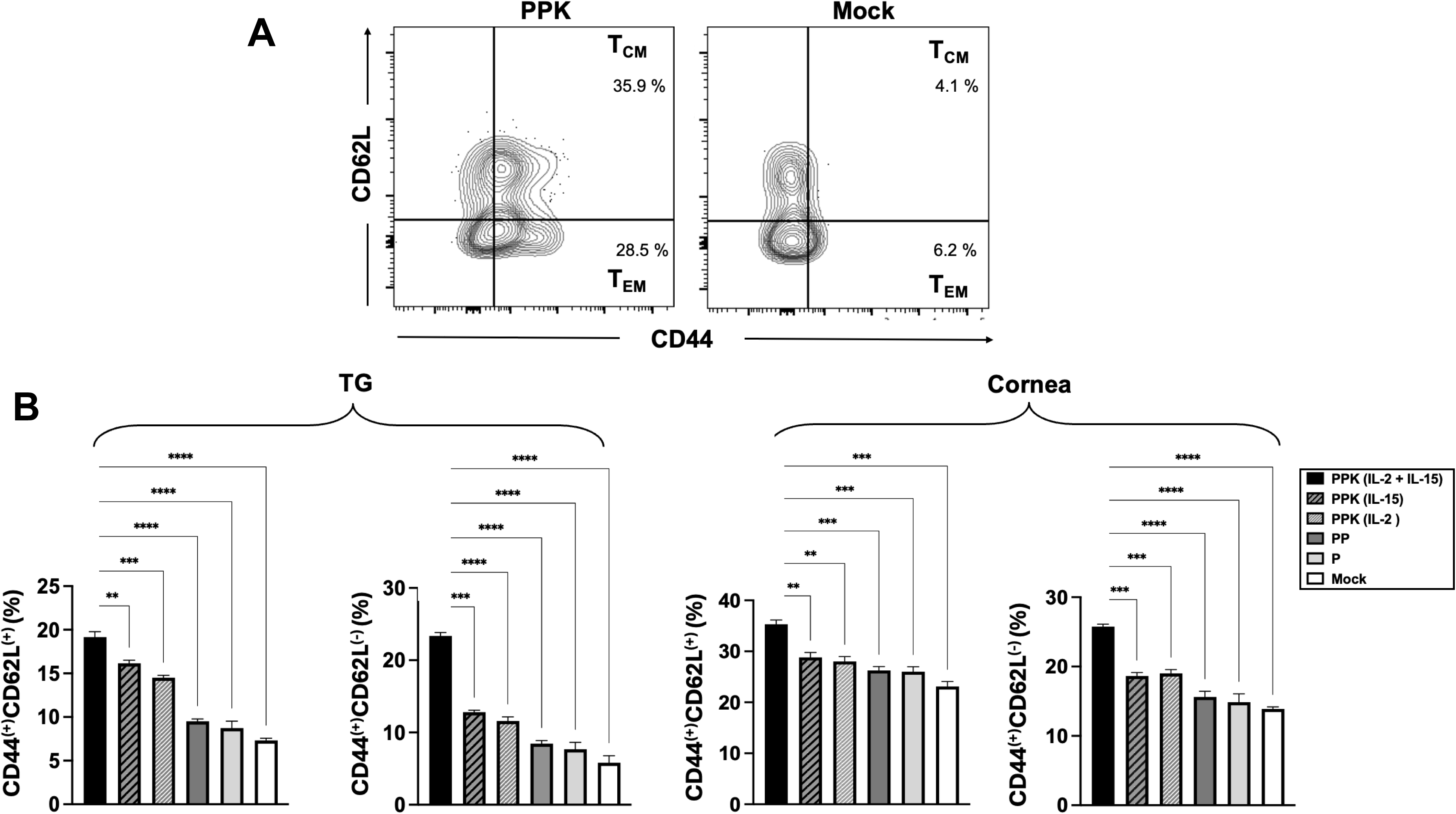
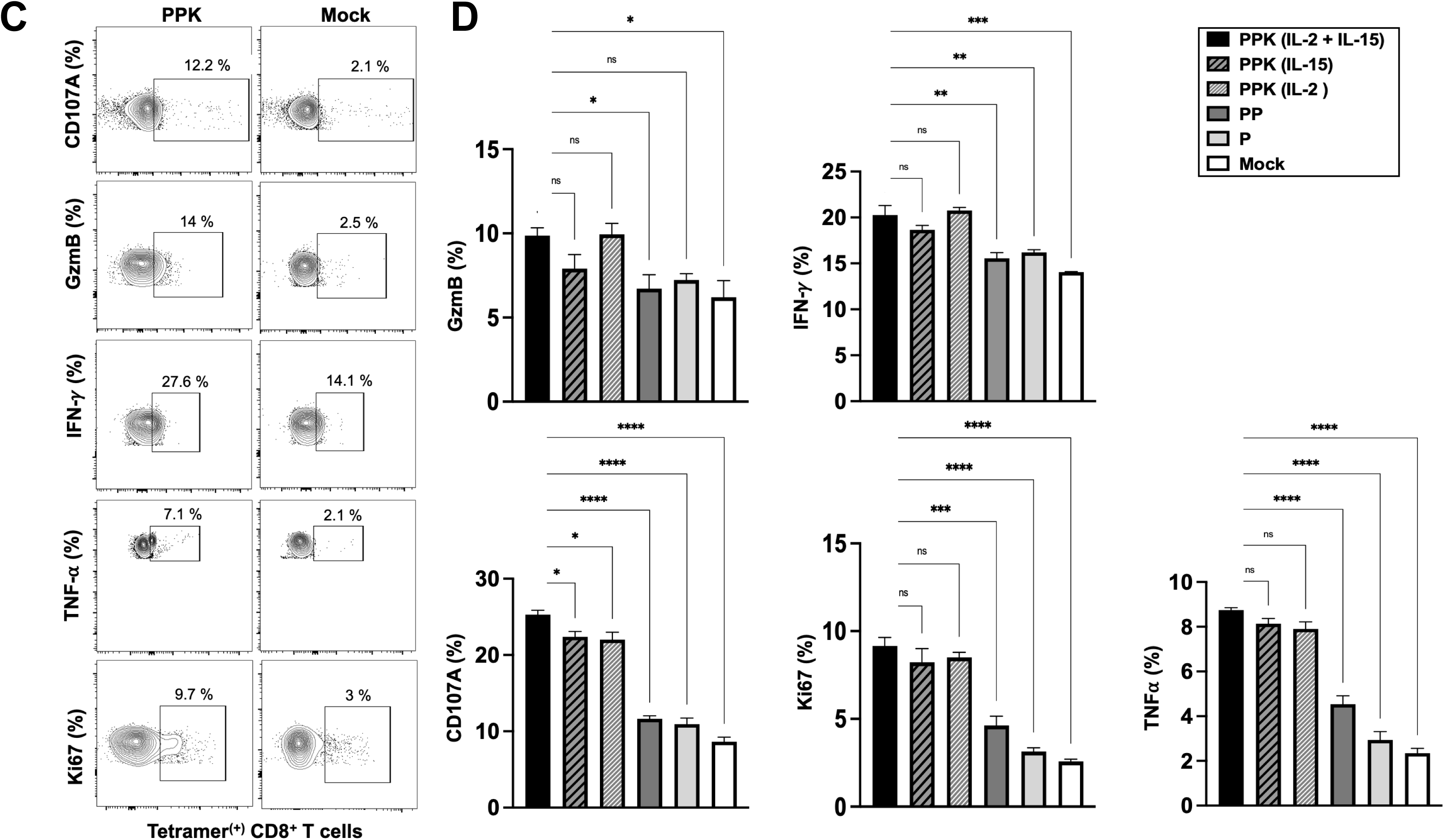
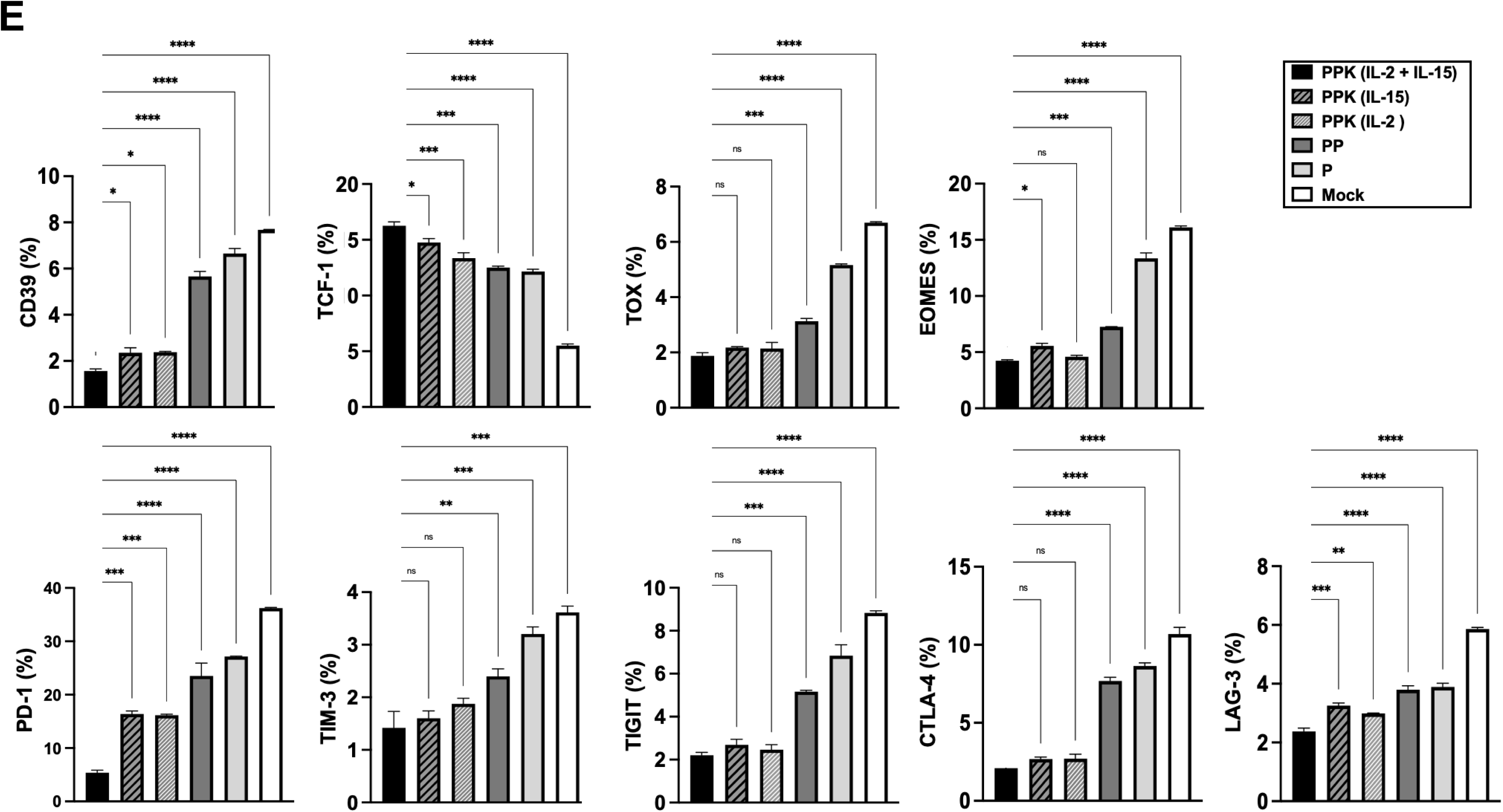
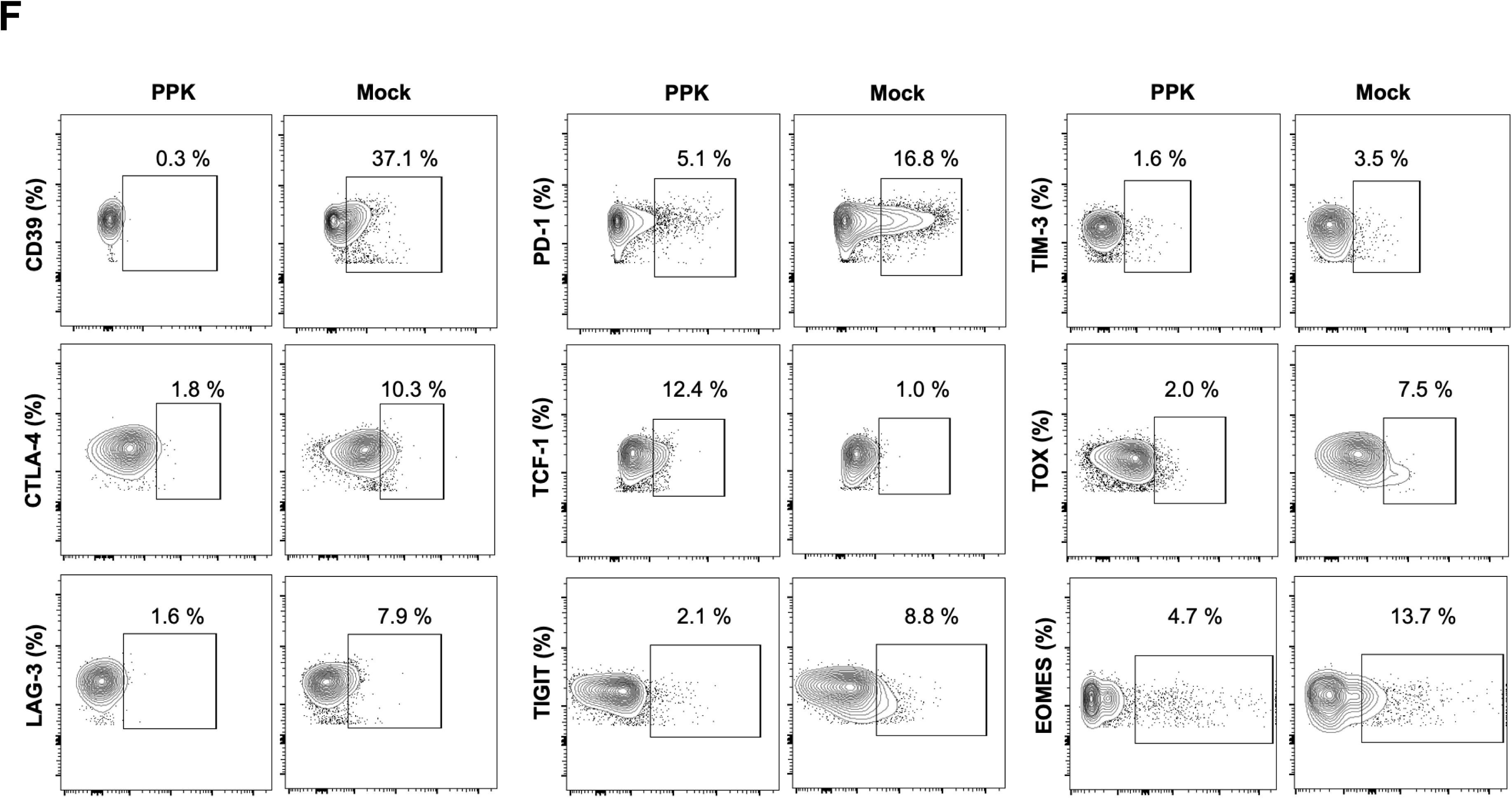
Prime/pull/keep T cell immunotherapy reduces HSV-1 viral loads in the HSV-1 infected HLA-A*0201/DR-B1 transgenic mice: A. Representative images of the trigeminal ganglia and corneal tissue immunoassayed for the presence of HSV-1 viral antigen when incubated with HSV-1 monoclonal antibody (Invitrogen-MA5-16285) followed by secondary staining with goat anti-mouse IgG (H+L), Alexa fluor™ 488 antibody (Invitrogen-A28175) in PPK treated and mock untreated HLA-A*0201/DRB1 Tg mice. Images were acquired at 20× magnification. B. Viral genome copy number was quantified by PCR from eye swabs collected from PPK-immunotherapy-treated and mock-untreated mice. A significant difference was defined as *\*P-value < 0.005*.

### 6. PPK Immunotherapy Enhances Polyfunctionality of HSV-1-Specific CD8^+^ T Cells, Boosting Cytokine Production, Cytotoxicity, and Proliferation in the Trigeminal Ganglia

To evaluate the functional profile of HSV-1-specific CD8^+^ T cells, the expression of IFN-γ, TNF-α, Granzyme B, CD107a, and Ki-67 was assessed by flow cytometry. PPK-containing IL-2 plus IL-15-treated mice exhibited significantly higher frequencies of IFN-γ (∼ 20 %) and TNF-α (∼9 %) producing cells compared with untreated controls, prime, or prime/pull group (**Figs. 4C** and **4D**), indicating enhanced antiviral effector function. These PPK-treated mice also showed increased expression of Granzyme B (∼10 %) and CD107a (∼9 %), markers associated with cytotoxic activity and degranulation, suggesting improved capacity to recognize and eliminate HSV-infected cells. The elevated expression of the proliferation marker Ki-67 (∼25 %) was also observed in the HSV-specific CD8^+^ T cells from PPK-treated mice, indicating enhanced expansion and maintenance of the antiviral T-cell response. These functional improvements were detected in the peripheral tissues of the TG, which significantly demonstrate that PPK immunotherapy increases the frequency of HSV-1-specific CD8^+^ T cells but also enhances their proliferative, cytokine-producing, degranulation, and cytotoxic capabilities. The generation of highly functional CD8^+^ T cells likely contributes to improved immune surveillance and control of recurrent HSV-1 infection.

### 7. PPK Immunotherapy Reduces CD8^+^ T Cell Exhaustion Markers and Promotes a TCF-1+ Self-Renewing Memory Phenotype in the Trigeminal Ganglia

To determine whether PPK immunotherapy influenced T cell exhaustion, the expression of multiple inhibitory receptors and transcriptional regulators associated with dysfunctional CD8^+^ T-cell responses was evaluated in HSV-1-specific tetramer-positive CD8^+^ T cells. Compared with untreated controls, PPK-containing homeostatic keep cytokines IL-2 plus IL-15-treated mice exhibited significantly lower frequencies of PD-1^+^, TIM-3^+^, TIGIT^+^, LAG-3^+^, CD39^+^, and CTLA-4^+^ CD8^+^ T cells in the TG, indicating reduced exhaustion and enhanced functional T cells (**Figs. 4E** and **4F)**. Analysis of exhaustion-associated transcription factors further demonstrated decreased expression of TOX and EOMES in HSV-1-specific CD8^+^ T cells from these PPK-treated mice, consistent with a less terminally exhausted phenotype. In contrast, expression of TCF-1, a transcription factor associated with self-renewal, memory formation, and sustained antiviral immunity, was significantly increased by ∼16% following PPK treatment. These findings indicate that PPK immunotherapy limits the development of T-cell exhaustion while promoting the maintenance of a memory and functional HSV-1-specific CD8^+^ T-cell population. This may contribute to improved immune surveillance and long-term control of recurrent HSV infection.

### 8. PPK Immunotherapy Elicits Robust HSV-1-Specific IgG Responses with a Th1-Biased Isotype Profile Complementing Mucosal Immunity

To determine whether PPK immunotherapy elicited HSV-1-specific humoral immunity, serum samples collected at the study endpoint were analyzed for antibodies against HSV-1 glycoprotein D (gD) and whole HSV-1 virions for HSV-1-specific antibodies, and antibody isotyping was performed using increasing serum dilution (between 1:20 and 1:312,500) by ELISA. PPK-containing both the cytokines IL-2 plus IL-15-treated mice exhibited significantly higher levels of HSV-1-gD or HSV-1-specific total IgG antibodies compared with untreated controls and pre-immune serum (**Fig. 5A**). All other groups, including PPK, PP, and prime, only showed significant levels of HSV-1-gD-specific total IgG antibodies, demonstrating the induction of a robust antigen-specific humoral immune response.

**Figure 5.**
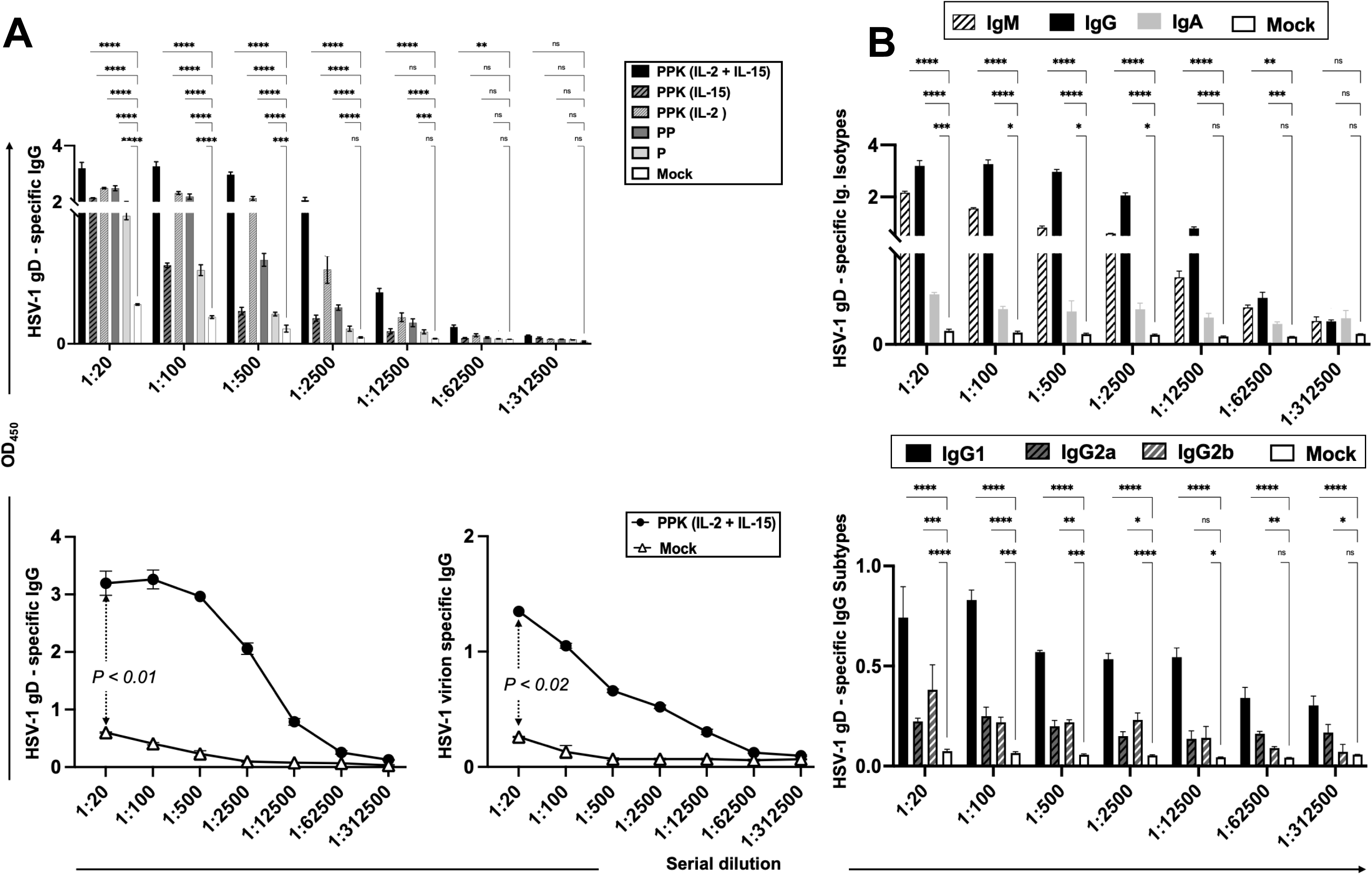
Prime/pull/keep T cell immunotherapy enhances HSV-1-specific CD8^+^ T cell responses in the trigeminal ganglion and cornea. Flow cytometric analysis of HSV-1-specific CD8^+^ T cells in the trigeminal ganglion and cornea. Higher frequencies of CD8^+^ T cells specific to HLA-A*0201-restricted HSV-1 epitopes (gD_53-61_, gB_183-191_, VP11/12_220-228_, VP13/14_504-512_, and UL9_196-204_) were detected by tetramer staining in PPK-treated mice compared with untreated mice. Experimental groups (P, PP, and PPK) and vaccination timelines are defined in Figure 1. Data are presented as mean ± SEM. Statistical significance was determined using a two-way ANOVA followed by multiple comparisons tests *(*P<0.05, ∗∗*P*<0.01, ∗∗∗P<0.001, ∗∗∗∗P<0.0001*; ns, not significant).

Antibody isotype analysis revealed that IgG was the predominant antibody detected, followed by IgM and then various IgG subtypes, such as IgG1, IgG2b, and IgG2a (**Fig. 5B**). IgA was also detected at lower levels (**Fig. 5B**). The strong IgM response indicates active antigen-specific B-cell activation, whereas the substantial IgG1 response reflects efficient class switching and the generation of a robust humoral immune response. The induction of IgG2a and IgG2b antibodies suggests Th1-associated responses and activation of antiviral effector mechanisms that may contribute to viral clearance and immune protection. In addition, detectable IgA responses indicate the development of mucosal-associated antibody responses. These findings demonstrate that PPK immunotherapy elicits a Th1-biased mixed immune response, which may complement cellular immune responses to enhance protection against recurrent HSV-1 infection and viral reactivation.

### 9. PPK Immunotherapy Restores Spatial Memory, Reduces Depression-Like Behavior, and Improves Social Function in HSV-1-Reactivated Mice

To determine whether PPK immunotherapy could mitigate neurological and behavioral deficits associated with recurrent HSV-1 infection, mice were evaluated by behavioral tests, including the Y-maze test, tail suspension test (TST), novel object recognition (NOR), novel object location (NOL), three-chamber sociability, and social novelty assays (**Fig. 6**). In the Y-maze test, PPK-treated mice exhibited significantly higher spontaneous alternation percentages compared with untreated HSV-1 reactivated controls, indicating improved spatial working memory (**Fig. 6A**). Consistent with these findings, mice receiving PPK immunotherapy demonstrated reduced immobility times in the tail suspension test, suggesting attenuation of HSV-1 associated depression-like behavior and improved neurobehavioral function (**Fig. 6B**). Cognitive performance was further evaluated using the NOR and NOL tests (**Figs. 6C** and **6D**). PPK-treated mice spent significantly more time exploring the novel object and the relocated object, respectively, compared with untreated controls, indicating improved recognition memory and hippocampal-dependent spatial memory. These findings suggest that PPK immunotherapy effectively preserves cognitive function following recurrent HSV-1 reactivation. Assessment of social behavior revealed that PPK-treated mice spent more time interacting with a novel mouse during the sociability assay and displayed a stronger preference for a novel versus a familiar mouse during the social novelty test (**Figs. 6E** and **6F**). These results indicate improved social interaction and social recognition memory compared with untreated HSV-1 reactivated mice. These findings demonstrate that PPK immunotherapy significantly ameliorates HSV-1-induced cognitive, behavioral, and social deficits, suggesting that enhanced antiviral immune control may protect against the long-term neurological consequences of recurrent HSV-1 infection.

**Figure 6.**
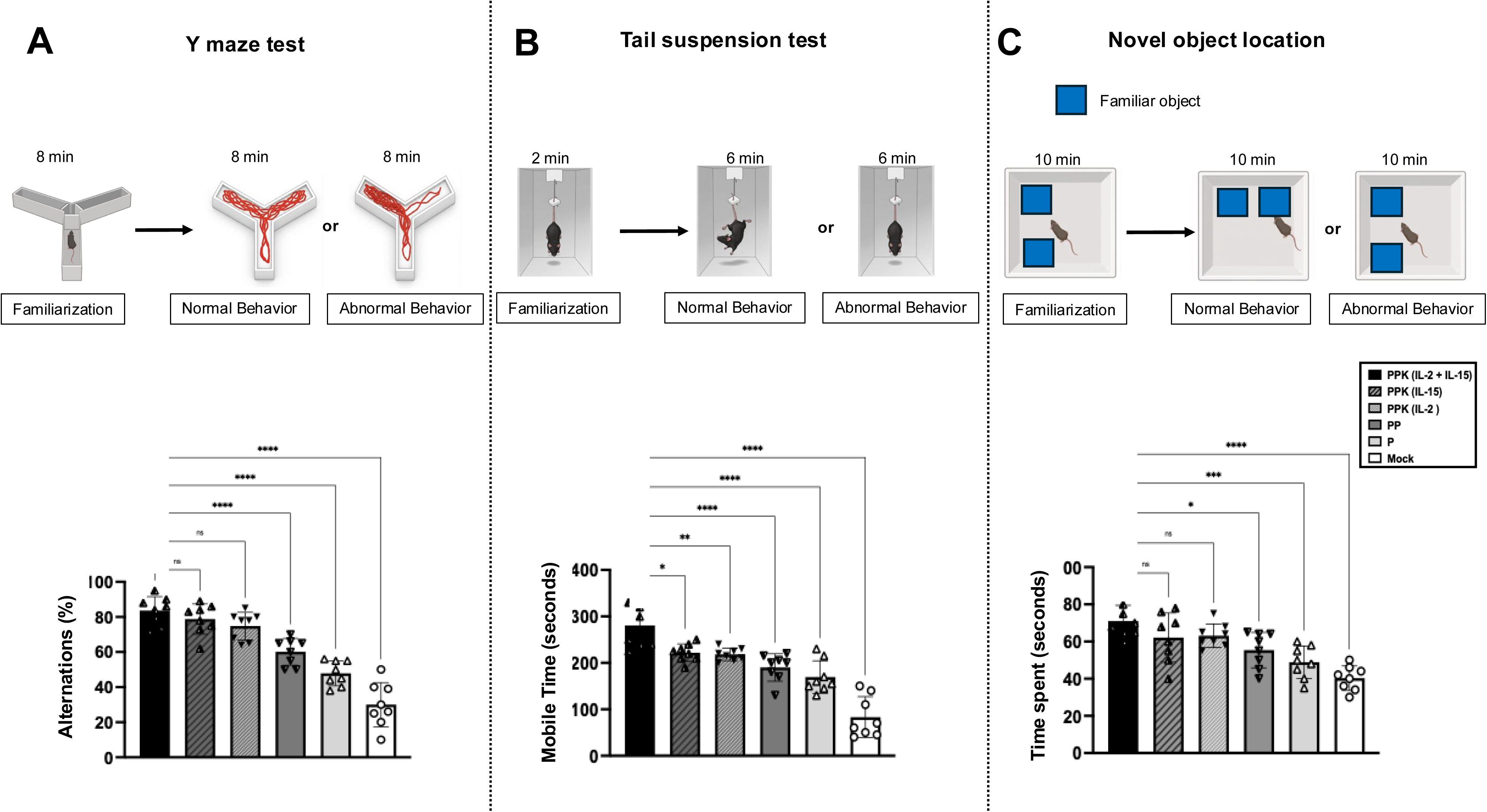
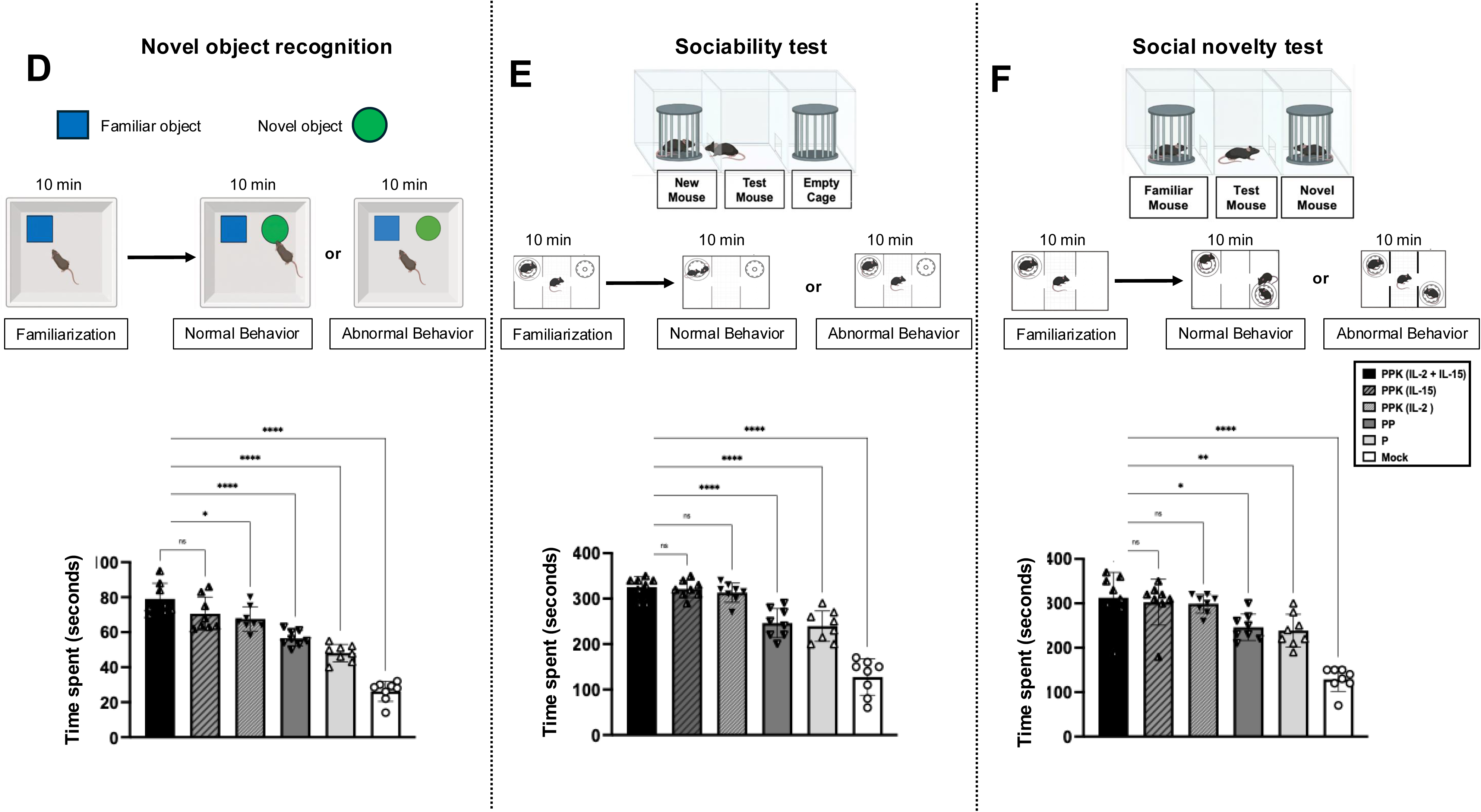
Prime/pull/keep T cell immunotherapy promotes tissue resident memory (T_RM_) responses in the trigeminal ganglion and cornea. Flow cytometric analysis of tissue resident memory (T_RM_) CD8+ T cell populations in trigeminal ganglion and cornea. Frequencies (%) of HSV-1-specific tissue-resident memory (T_RM_) markers were analyzed on gated CD8^+^ T cells isolated from the trigeminal ganglia (TG) and cornea of PPK-treated and untreated mice. Expression of CD49a, CD69, and CD103 was assessed within tetramer-positive CD8+ T cells specific for the gB_183-191_ epitopes. Data are presented as the mean ± SEM. Statistical significance between groups was determined using one-way ANOVA followed by multiple comparisons tests (*∗P<0.05, ∗∗P<0.01, ∗∗∗P<0.001, ∗∗∗∗P<0.0001*; ns, not significant).

Thus, our findings (i) demonstrate that HSV-1-specific peptide epitopes combined with CXCL11 chemokine delivery and IL-2/IL-15 cytokine support significantly reduce ocular HSV-1 viral burden and attenuate the severity of recurrent herpes corneal infection in mice; (ii) indicate that the enhanced protection is associated with improved quantity and functional quality of HSV-1-specific CD8⁺ T cells infiltrating the cornea and trigeminal ganglia under a prime/pull/keep strategy, including a reduced expression of exhaustion markers. ELISA-based antibody profiling and isotyping; (iii) ELISA and isotyping further show that vaccination induces a favorable humoral response characterized by a balanced immunoglobulin isotype distribution consistent with a mixed Th1/Th2 immune profile; and (iv) demonstrate that vaccinated mice exhibit improved behavioral outcomes in disease-associated assays, consistent with reduced ocular pathology and neuroinflammation. These results significantly support the HLA-A*02:01/DRB1 transgenic mice as a relevant preclinical model of ocular herpes infection and disease.

## DISCUSSION

Recurrent ocular herpes remains a substantial global health burden. HSV-1 ocular infection produces a broad spectrum of clinical disease, ranging from relatively mild blepharitis and conjunctivitis to dendritic epithelial keratitis, disciform stromal edema, and ultimately blinding herpes stromal keratitis, which remains a leading infectious cause of corneal blindness in the developed world (11). While current antiviral therapies, including acyclovir and its derivatives, reduce disease severity, they do not eliminate the virus that reactivates from tissues or reliably prevent recurrent ocular disease and progression to corneal blindness. Despite decades of clinical development, no HSV therapeutic vaccine has yet succeeded in large-scale clinical trials (63, 64). A central obstacle is HSV-1’s capacity to establish lifelong latency in sensory neurons, particularly the ganglia, from which it periodically and sporadically reactivates to cause recurrent herpetic disease (65–67). Compounding this, HSV-1 has evolved extensive mechanisms to suppress host innate and adaptive immunity, including tissue-resident CD4^+^ and CD8^+^ T cells, complicating efforts to achieve durable vaccine-induced protection (16, 68). Thus, traditional protein-based vaccines, even when mixed with strong immuno-adjuvants, are mostly unsuccessful in inducing antiviral tissue-resident CD4^+^ and CD8^+^ T cells and have shown limited protective efficiency against recurrent HSV infection and disease (13, 69, 70). These protein-based vaccines induce neutralizing antibody responses, but they fail to generate robust and durable tissue-resident memory (T_RM_) CD4^+^ and CD8^+^ T cell responses, which are critical for controlling viral reactivation at neuronal sites (15, 71). This underscores the need for a safe, effective, T-cell-targeted vaccine or immunotherapies to boost antiviral CD4^+^ and CD8^+^ T_RM_ cells that will then eliminate the tissue-hiding virus reservoir, the root of the disease (72). In the present study, we present a novel PPK T-cell-based immunotherapy, built around HSV-1 CD8^+^ and CD4^+^ T-cell immunodominant epitopes presented by human HLA molecules and preferentially recognized by naturally protected ASYMP individuals. This PPK T-cell-based immunotherapy boosted antiviral CD4^+^ and CD8^+^ T_RM_ cells and reduced virus reactivation, shedding, recurrent corneal disease, and alleviated cognitive, behavioral, and social deficits in a ‘humanized’ mouse model of UVB-induced reactivation and recurrent ocular herpetic disease.

Tissue-resident memory T cells (T_RM_) reside primarily in peripheral tissues, where they provide rapid and robust immunosurveillance against invading pathogens, malignant cells, and other inflammatory or autoimmune diseases (73). T_RM_ cells: (*i*) provide localized, site-specific protection in peripheral tissues, such as the mucosal surface (74); (*ii*) crosstalk with dendritic cells (DCs), enhance local immune response, and support anti-tumor immunity (75); (*iii*) are more effective than circulating memory T cells in maintaining long-term immunity (27); (*vi*) play a critical role in preserving immune homeostasis and tissue integrity; and (*v*) have broader implications for human health, including protection against recurrent viral infections such as herpes (76). In the present study, we demonstrate that a Tissue-targeted T cell immunotherapy elicits robust and durable mucosal and tissue-resident memory T_RM_ responses, holds the potential to reduce viral recurrence, limit corneal inflammation, and inhibit herpes-related blindness (46, 77–79).

The multi-epitope PPK construct represents a rational advance over single-antigen HSV-1 vaccine approaches by concentrating the immune response on epitopes empirically validated in asymptomatic individuals, a population of herpes patients whose natural, HSV-1-specific T cell responses correlate with protection from recurrent disease. By combining HLA-restricted human CD4^+^ and CD8^+^ immunodominant epitopes from HSV-1 structural (gD, gB), tegument (VP11/12, VP13/14), and replication-associated (UL9) proteins, this design broadens antigenic coverage across distinct stages of the viral life cycle, potentially reducing the likelihood of immune escape compared to single-glycoprotein candidates such as gD2, which have previously failed in clinical trials (63, 64). Selection of epitopes based on high HLA-A*0201/DR-B1 binding affinity further supports translational relevance, as these haplotypes are common in the general population and predictive of efficient antigen presentation. The use of AAV8 under a neuron-specific CamKIIα promoter is a deliberate strategy to achieve local, sustained epitope expression within trigeminal ganglion neurons, the anatomical site of HSV-1 latency, rather than relying on transient peripheral immunization, aligning with the “prime/pull/keep” logic of first priming systemic T cells, then pulling and retaining them at the site of latent infection. Co-expression of CXCL11 T-cell attracting chemokine, to recruit T cells, and IL-2/IL-15 survival cytokines to sustain tissue-resident memory CD4^+^ and CD8^+^ T cell populations addresses a well-recognized limitation of prior therapeutic vaccine efforts: even when antigen-specific T cells are generated, they often fail to establish durable residency within latently infected ganglia. Together, this construct design directly targets the immunological bottlenecks, narrow antigen breadth, poor CD4^+^ and CD8^+^ T cell trafficking, and lack of long-term retention that have limited the success of earlier HSV-1 vaccine candidates.

Several cytokines and chemokines are reported to play critical roles in modulating T cell recruitment (80, 81). CXCR3-expressing T cells migrate in response to gradients of interferon-inducible C–X–C chemokine ligands, including CXCL9 (MIG), CXCL10 (IP-10), and CXCL11 (I-TAC) (41). The role of the CXCL9/CXCL10/CXCL11-CXCR3 axis in the pathophysiology of Th1 predominant diseases, such as viral infection, has been reported by several investigators (82). Our lab has demonstrated the role of CXCL11 in recruiting effector T cells during acute infection as well as infiltration of CXCR3⁺ T cells into mucosal tissues and retention of virus-specific tissue-resident memory T cells (T_RM_) in genital and respiratory mucosa of Herpes simplex virus (HSV) and SARS-CoV-2 infections, respectively (46, 79, 83). The role of CXCL11 as a T cell modulator has also been explored in multiple cancers (82, 84). The secretion of CXCL11 increases the infiltration of CD8^+^ T cells into the tumor microenvironment and suppresses angiogenesis (32, 85). These findings highlight the potential of targeting the CXCL11– CXCR3 axis to enhance mucosal vaccine strategies by improving local immune protection. T cell retaining cytokines, IL-2 and IL-15, are critical for the generation, maintenance, and functional enhancement of memory T cells (33). We have recently reported that co-delivery of IL-2 or IL-15 as keep has resulted in increased tissue-resident memory T cell formation in Herpes ocular infection in the rabbit model (46). Previous studies have shown that IL-2 promotes the expansion of CD4^+^ T_RM_ populations and enhances protective efficacy against invasive nontyphoidal salmonellosis (86). Similarly, IL-15 has been reported to enhance lung resident memory T cells, further underscoring its importance in establishing durable mucosal immunity (87). Both cytokines play a crucial role in anti-tumoral immunity by promoting and maintaining tumor resident memory T cells (88).

Our findings demonstrate that therapeutic protection against recurrent ocular herpes requires all three components of the PPK strategy acting in concert, with the “keep” cytokines (IL-2/IL-15) emerging as the decisive determinant of durable efficacy. The UVB-induced reactivation model in HLA-A*0201/DR1 transgenic mice is a well-validated system for this purpose, as it reliably reproduces environmentally triggered HSV-1 reactivation and recurrent herpetic stromal keratitis within a “humanized” antigen-presentation background (89–97), allowing preclinical evaluation of human HLA-restricted T cell epitope vaccines before progression to clinical trials. The graded improvement observed from prime alone, to prime/pull, to full PPK treatment mirrors the established immunological rationale that generating antigen-specific T cells (prime) and recruiting them *via* the T cell-attracting chemokine signaling (pull, CXCL11) are necessary but insufficient steps unless those cells can also be retained by survival cytokines, as a stable tissue-resident memory population within the TG and cornea, the anatomical reservoir of HSV-1 latency. The convergence of improved survival, preserved corneal epithelial integrity on fluorescein staining, reduced histopathological damage, and lower viral antigen and genome burden in the PPK/cytokine groups indicates that the survival cytokines IL-2/IL-15-supported T cell retention translates directly into enhanced local viral control, consistent with prior work showing that cytokine-supported tissue-resident memory CD4^+^ and CD8^+^ T_RM_ cells in the TG are critical for suppressing HSV-1 reactivation and shedding. Together, these results support a model in which durable protection against recurrent ocular herpes is governed less by the magnitude of the initial T cell response than by the ability to sustain that response at the tissue site of latency, as we previously discussed (98).

While PPK1 (IL-2) and PPK2 (IL-15) monotherapies provided substantial protection over prime/pull alone, PPK3 (IL-2+IL-15) consistently demonstrated superior efficacy across both tissue-resident memory retention and behavioral recovery. Direct pairwise comparison showed PPK3 (IL-2+IL-15) significantly outperformed PPK1 (IL-2 alone) and PPK2 (IL-15 alone), while PPK1 and PPK2 did not differ significantly from each other in most assays. This synergy likely reflects the distinct, complementary homeostatic signals provided by the two cytokines. IL-2 drives robust initial expansion and effector differentiation of antigen-specific T cells, whereas local IL-15 signaling is critical for the long-term survival and homeostatic proliferation of CD4⁺and CD8⁺ T_RM_ cells without inducing exhaustion. By delivering both signals simultaneously *via* the AAV8 platform, PPK3 optimizes both the initial deployment and the local retention of protective T_RM_ populations within the cornea and trigeminal ganglia. Thus, our tissue-targeted topical ocular PPK T cell immunotherapy blocks the HSV-1 latency-reactivation cycle, promotes a protective immune environment by increasing CXCL11, IL-2, and IL-15 expression, generates robust CD49a⁺CD69⁺CD103⁺ tissue-resident memory T cells, enriches CD44⁺CD62L^+^, CD44^+^CD62L**^−^** memory T cell subsets, enhances TCF-1 expression, and reduces the expression of exhaustion-associated markers.

The upregulation of CD69, CD103, and CD49a on tetramer-positive CD8^+^ T cells confirms that the PPK immunotherapy drives HSV-1-specific T cells toward a bona fide tissue-resident memory T_RM_ phenotype rather than simply increasing their number, a distinction that matters functionally since CD69 restrains tissue egress by antagonizing S1PR1-mediated recirculation, while CD103 mediates E-cadherin–dependent epithelial anchoring and CD49a supports long-term persistence and cytotoxic poise at barrier sites, with CD69 driving early retention and CD103 sustaining long-term lodgment of T_RM_ cells in tissue. This T cell signature is consistent with prior work in genital HSV-2 infection showing that tissue-resident CD8^+^ T_RM_ cells bearing these same markers serve as first-responders that intercept virus at the earliest stage of reactivation, aborting clinical recurrence before lesions can form (99–101). The compartmentalized enrichment of T_RM_ cell markers in the cornea and TG, but not the spleen, together with the corresponding histological infiltration of CD4^+^, CD8^+^, and CD103^+^ cells specifically in chemokine/cytokine-supplemented PPK groups, indicates that CXCL11-mediated recruitment and IL-2/IL-15-mediated retention act sequentially and synergistically to convert a transient effector response into a stable, site-anchored T_RM_ cell population. Collectively, these results support a model in which the PPK strategy’s therapeutic benefit is mechanistically explained by its capacity to establish durable T_RM_ cell immunity precisely at the anatomical interface of HSV-1 latency and reactivation, providing continuous local immunosurveillance that limits viral spread and tissue pathology.

The phenotypic characterization of T cell populations further supports the generation of durable tissue-directed immunity following PPK T cell immunotherapy. Increased frequencies of CD49a⁺, CD69⁺, and CD103⁺ T cells are consistent with the establishment of tissue-resident memory T cells (T_RM_) (102–104). CD69 functions as a tissue-retention molecule; CD103 promotes adhesion to epithelial cells through interactions with E-cadherin, facilitating long-term residency at peripheral sites (104). In addition, CD49a (integrin α1) has been associated with highly functional resident memory populations exhibiting enhanced cytotoxic potential and increased production of antiviral effector molecules, including IFN-γ and granzyme B (103). Analysis of memory T cell subsets revealed increased frequencies of CD44⁺CD62L⁻ effector memory T cells (T_EM_), which rapidly migrate to peripheral tissues and exert immediate effector functions upon antigen encounter (78). In contrast, CD44⁺CD62L⁺ central memory T cells (T_CM_) are characterized by enhanced proliferative capacity and long-term persistence (105). The expansion of both T_EM_ and T_RM_ populations suggests the development of a robust memory response capable of providing rapid local protection while maintaining long-term immunological surveillance. These findings also correlate with earlier studies (78). The importance of CXCR3-dependent chemokine signaling in antiviral immunity using the CXCL10/CXCR3 and CXCL11/CXCR3 axis has been investigated earlier, which showed HSV-1-specific T_EM_, T_CM,_ and T_RM_ cells within infected tissues, thereby enhancing recruitment of activated CXCR3-expressing T cells into peripheral tissues and protection against recurrent herpesvirus infection (79, 106). Once localized within the tissue microenvironment, these cells may differentiate into resident memory populations under the influence of cytokines such as IL-2 and IL-15, which support T cell expansion, survival, memory maintenance and establish protective T_RM_ populations (73).

Furthermore, the infected mice treated with our PPK T cell immunotherapy exhibited a less exhausted T cell phenotype. Chronic antigen stimulation typically induces progressive expression of inhibitory receptors, including PD-1, TIM-3, LAG-3, TIGIT, CTLA-4, and CD39 as well as exhaustion-associated transcriptional regulators such as TOX and EOMES, leading to functional impairment of T cells (47, 107). The reduced expression of these exhaustion markers suggests preservation of T cell function, which may enhance the ability of memory populations to respond effectively upon antigen re-exposure. In contrast, elevated expression of multiple inhibitory receptors is generally associated with diminished cytokine production, impaired proliferative capacity, and reduced cytotoxic activity. Increased expression of TCF-1, a transcription factor that plays a central role in T cell development and the generation of long-lived memory (108).TCF-1 functions as a hallmark of stem-like memory and progenitor exhausted T cells that retain proliferative potential and contribute to durable immune responses and support long-term immune surveillance and recall responses.

The predominance of gD- and virion-specific IgG, alongside detectable IgG2a and IgG2b subclasses, indicates that PPK immunotherapy elicits a Th1-polarized humoral response, consistent with the well-established link between IFN-γ signaling and class switching to IgG2a in mice, since HSV-1 is a strong inducer of Th1-type immune responses in which IFN-γ and CD4^+^ T cells producing it drive isotype switching to IgG2a during infection. This Th1-skewed antibody profile is functionally significant, as IgG2a is generally considered the most effective isotype for antiviral clearance through Fc-mediated effector mechanisms such as opsonization and complement fixation, in contrast to the largely non-neutralizing, Th2-associated IgG1 response typical of poorly protective vaccines. The concordance between this humoral Th1 bias and the cellular IFN-γ/granzyme B/TCF-1 profile described above suggests that PPK immunotherapy engages a coordinated, IFN-γ-driven axis linking CD4^+^ T cell help, B cell isotype switching, and CD8^+^ T cell effector function, rather than inducing cellular and humoral immunity independently. The presence of mucosal IgA further suggests that PPK immunotherapy also seeds antibody responses capable of acting at mucosal surfaces relevant to ocular HSV-1 shedding and reinfection. Collectively, these findings support the interpretation that PPK immunotherapy-induced protection against recurrent ocular herpes reflects an integrated Th1-biased cellular and humoral response rather than a single dominant effector arm.

The amelioration of spatial working memory, depression-like behavior, recognition memory, and social behavior in PPK-treated mice suggests that improved local viral control at the TG and cornea translates into protection against the broader neurological consequences of recurrent HSV-1 reactivation. This is consistent with prior work showing that repeated cycles of HSV-1 reactivation drive viral spread into the hippocampus and cortex, triggering neuroinflammation, astrogliosis, and progressive, reactivation-dependent cognitive deficits that become irreversible with cumulative reactivation cycles. More broadly, HSV-1-associated neuroinflammatory signaling, independent of overt encephalitis, has been linked to cognitive, anxiety-like, and social/motor impairments in mouse models, supporting the interpretation that chronic immune activation and viral persistence, rather than acute disease alone, underlie these behavioral deficits. By restricting viral replication and reactivation at the ganglionic reservoir through durable, non-exhausted T_RM_ cell immunity, PPK immunotherapy likely limits the downstream neuroinflammatory cascade responsible for these deficits, rather than acting directly on the CNS. Taken together with the virological, cellular, humoral, and pathological findings, these behavioral outcomes provide functional, whole-organism evidence that PPK immunotherapy-induced immune protection is not confined to the ocular surface but extends to preserving neurological and psychosocial function, and they reinforce the HLA-A*02:01/DRB1 transgenic, UVB-reactivation mouse model as a translationally relevant platform for evaluating next-generation therapeutic vaccines against recurrent ocular herpes.

The ultraviolet B (UVB) induced herpes recurrence murine model closely mimics environmentally triggered reactivation of HSV-1 in humans (59). After primary ocular infection, mice develop recurrent herpetic stromal keratitis, blepharitis, and viral shedding, paralleling the clinical symptoms of human ocular herpes (8). Thus, this small-breeding, cost-effective, and immunologically relevant mouse model provides critical insights for the development of effective therapies by enabling the investigation of the protective role of CD8^+^ T cells against primary herpes simplex virus infection (109). While human HLA is highly polymorphic, HLA-A*0201 is one of the most widely distributed alleles globally. Using this specific humanized transgenic model ensures that the identified protective epitopes carry direct translational validity for a large percentage of (∼ 50 %) human clinical populations, bridging the gap between mouse models and human clinical trials (110). In the present study, we utilized an HLA-A*0201/DR-B1 transgenic mouse model of ocular herpes to investigate the therapeutic potential of our multiepitope construct.

HSV glycoprotein B (gB), glycoprotein D (gD), viral tegument proteins (VP11/12 and VP13/14), and the DNA replication-binding helicase (UL9) are the major HSV antigens and have been reported to generate protective immunity in both animal models and humans (46, 111). Symptomatic patients often exhibit reactivation of latent HSV-1, leading to the expansion of “pathogenic” HSV-1-specific CD4⁺/CD8⁺ T cells that contribute to recurrent herpetic disease (76, 112). In contrast, asymptomatic individuals harbor the virus without experiencing recurrent symptoms, suggesting a protective and HSV-specific immune response (58). Our several reports on ASYMP HSV-antigen-specific therapeutic vaccines, including full-length, partial, and epitope-based therapeutic vaccines, have demonstrated protective efficacy (44, 46, 78, 79). The epitope-based vaccine encoding gD has shown reduced spontaneous reactivation by 4-to 5-fold, which is significantly higher in rabbits (∼57%) and humans (58%) reported using partial or whole gD protein (113, 114). The ASYMP CD8^+^ T cell epitopes-based peptide vaccine using HSV-1 membrane glycoprotein C (UL44_400–408_), the DNA replication binding helicase (UL9_196–204_), and the tegument protein (UL25_572–580_) has demonstrated protective efficacy against ocular herpesvirus infection (96). To overcome the limitations of current HSV vaccines and induce long-term protective immunity, we developed a multi-epitope T-cell-based immunotherapy construct incorporating both CD8⁺ and CD4⁺ T cell epitopes derived from HSV-1 glycoprotein B (gB), glycoprotein D (gD), viral tegument proteins (VP11/12 and VP13/14), and the DNA-binding helicase UL9. These epitopes were selected based on immune profiling of naturally protected asymptomatic (ASYMP) individuals, who exhibit a higher frequency of HSV-1-specific effector memory CD8⁺ T cells (45, 115).

Non-replicating recombinant adeno-associated viral (AAV) vectors have been widely explored for tissue-specific delivery of therapeutic genes (34). Currently, more than 70% of gene delivery vectors used in clinical trials are viral-based, highlighting their efficiency and versatility (35). Among these, AAV serotype 8 (AAV8) has been extensively studied and utilized in gene therapy research for a broad range of diseases, including genetic disorders, cancers, autoimmune conditions, and viral infections (34, 35, 77, 116). These neurotropic AAV8 vectors exhibit broad tissue tropism, efficiently transducing a variety of tissues, including the brain, liver, and lungs. Compared to other AAV serotypes, AAV8 delivers approximately 2.5-fold higher levels of vector DNA and enhances 10-to 100-fold higher expression of target genes, making it a highly effective vector for therapeutic applications (35). AAV8 vector achieves stable transduction with near 100% efficiency, underscoring its safety and potency for tissue-targeted gene therapy (35). To deliver T cell-attracting chemokines and T cell-retaining cytokines, we have employed an AAV8 vector in our prime/pull/keep (PPK) T cell immunotherapy studies. Due to the humanized HLA-restricted epitopes and a P2A self-cleaving linker, all three functional arms (prime/pull/keep) are included in a single stable open reading frame within the packaging limit of the AAV8 vector. Our recent work demonstrated the effective use of the AAV8 vector for tissue-specific delivery in both prime/pull and prime/pull/keep strategies targeting genital herpes, recurrent ocular herpes, and SARS-CoV-2 infection (46, 79, 83).

In conclusion, our study strongly supports PPK immunotherapy for inducing durable, functional mucosal T cell immunity against herpes related ocular infection. This strategy promoted recruitment and long-term maintenance of tissue-resident CD84⁺ and CD8⁺ T cells, leading to improved control of ocular recurrent HSV-1 infection and attenuation of disease pathology. These findings reinforce the value of this platform for inducing durable, localized antiviral immunity at the site of infection.

**Figure 7.** Prime/pull/keep T cell immunotherapy promotes infiltration of tissue residentmemory (T_RM_) cells: (**A**) Representative immunofluorescence images of the trigeminal ganglia collected at the endpoint from PPK T cell immunotherapy-treated and mock untreated mice. Tissue sections were stained using antibodies: Alexa Fluor® 488 anti-mouse CD4 antibody (Biolegend-100423), PerCP anti-mouse CD8a antibody (Biolegend-100732), PerCP/Cyanine5.5 anti-mouse CD103 antibody (Biolegend-121416); markers associated with tissue-resident memory (T_RM_) cells. Images were taken at 20× magnification. (**B**) Representative immunofluorescence images of cornea tissue collected at the endpoint from PPK immunotherapy-treated and mock untreated mice. Tissue sections were stained using antibodies: Alexa Fluor® 488 anti-mouse CD4 antibody (Biolegend-100423), PerCP anti-mouse CD8a antibody (Biolegend-100732), PerCP/Cyanine5.5 anti-mouse CD103 antibody (Biolegend-121416); markers associated with tissue-resident memory (T_RM_) cells. Images were taken at 20× magnification.

**Figure 8.** Prime/pull/keep T cell immunotherapy enhances central (T_CM_) and effector memory (T_EM_) CD8^+^ T cell responses in the trigeminal ganglion and cornea. Flow cytometric analysis of central memory (T_CM_; CD44^+^CD62L^+^) and effector memory (T_EM_; CD44^+^CD62L^−^) CD8^+^ T cell subsets in the trigeminal ganglia (TG) and cornea. Frequencies (%) of central memory (T_CM_ CD44^+^CD62L^+^) and effector memory (T_EM_, CD44^+^CD62L^−^) cells were assessed within gB_183-191_ tetramer-positive CD8^+^ T cells from PPK immunotherapy-treated mice compared to untreated control mice. Experimental groups (P, PP, and PPK) and vaccination timelines are defined in Figure 1. Data are presented as the mean ± SEM. Statistical significance between the experimental groups was evaluated using one-way ANOVA followed by appropriate multiple comparisons tests (*∗P<0.05, ∗∗P<0.01, ∗∗∗P<0.001, ∗∗∗∗P<0.0001)*.

**Figure 9.** Prime/pull/keep T cell immunotherapy enhances functional T cell markers in the trigeminal ganglion. **F**requencies (%) of CD8^+^ T cells expressing cytotoxic and effector function markers, including granzyme B (GzmB), interferon-γ (IFN-γ), tumor necrosis factor-α (TNF-α), CD107a, and the proliferation marker Ki67, were assessed in trigeminal ganglia. Experimental groups (P, PP, and PPK) and vaccination timelines are defined in Figure 1. Data are presented as the mean ± SEM. Statistical significance between the experimental groups was determined using a two-way ANOVA followed by appropriate multiple comparisons tests (*∗P<0.05, ∗∗P<0.01*, *∗∗∗P<0.001, ∗∗∗∗P<0.0001*; ns, not significant).

**Figure 10.** Prime/pull/keep T cell immunotherapy reverses T cell exhaustion phenotypes in the trigeminal ganglion. Flow cytometric analysis of exhaustion marker expression in the trigeminal ganglion. Frequencies (%) of exhaustion-associated markers including TCF-1, TO, EOMES, PD-1, LAG-3, TIM3, TIGIT, CTLA-4, CD39 were assessed on CD8^+^ T cells from PPK immunotherapy treated and untreated mice. Experimental groups (P, PP, and PPK) and vaccination timelines are defined in Figure 1. Data are presented as the mean ± SEM. Statistical significance was evaluated using two-way ANOVA followed by appropriate multiple comparisons tests *(∗P<0.05, ∗∗P<0.01, ∗∗∗P<0.001, ∗∗∗∗P<0.0001*; ns, not significant).

**Figure 11.** Prime/pull/keep T cell immunotherapy induces humoral immune responses in HSV-1-infected HLA-A*0201/DR-B1 transgenic mice. (**A**) HSV-1-gD-specific serum antibody titers were measured in the PPK immunotherapy treated, untreated control groups across serial serum dilution points ranging from 1/20 to 1/312,500. (**B**) HSV-1-gD-specific antibody titer responses were assessed in mice receiving PPK (IL-2 + IL-15) treatment and untreated mice across serial serum dilution points ranging from 1/20 to 1/312,500. (**C**) HSV-1 virion-specific antibody titers were measured in the PPK (IL-2 + IL-15) treated and untreated mice across serial serum dilution points ranging from 1/20 to 1/312,500. (**D**) Serum immunoglobulin isotypes-IgM, IgA, and IgG profile were evaluated against HSV-1-gD and compared between PPK (IL-2+IL-15) immunotherapy treated with untreated mice. (**E**) IgG subtypes (IgG1, IgG2a, or IgG2b specific to HSV-1 gD were analyzed and compared between PPK (IL-2+il-IL-15) immunotherapy treated with untreated mice. Experimental groups (P, PP, and PPK) and vaccination timelines are defined in Figure 1. Data are presented as the mean ± SEM. Statistical significance across the experimental groups and individual dilution points was determined using two-way ANOVA followed by appropriate multiple comparisons tests (*∗P<0.05, ∗∗P<0.01, ∗∗∗P<0.001, ∗∗∗∗P<0.0001*; ns, not significant).

**Figure 12.** Prime/pull/keep T cell immunotherapy prevents behavioral deficits in treated mice. Behavioral performance was evaluated using the following tests. (**A**) Y-maze test: Spatial working memory was assessed by measuring spontaneous alternation behavior or the percentage of alternation entries. (**B**) Tail suspension test: Depressive behavior was evaluated by measuring the duration of immobility during the test period. (**C**) Novel object recognition (NOR) test: Recognition memory was assessed by determining the time spent exploring the novel versus familiar object. (**D**) Novel object Location test: Memory performance was evaluated by measuring the time spent exploring the object in the novel location compared with the object remaining in the familiar location (**E**) Three-chamber sociability test: Social preference was assessed by comparing the time spent interacting with a novel mouse versus an empty chamber. (**F**) Social novelty test: Social recognition memory was evaluated by measuring the preference for interaction with a novel stranger mouse compared with a familiar mouse. Data are presented as mean ± SEM (*n* = 8 animals per group). Experimental groups (P, PP, and PPK) and vaccination timelines are defined in Figure 1. Statistical significance between the experimental groups was evaluated using one-way ANOVA followed by multiple comparisons tests *(∗P<0.05, ∗∗P<0.01, ∗∗∗P<0.001, ∗∗∗∗P<0.0001*; ns, not significant).

## ACKOWLEDGEMENTS

This work is supported by Public Health Service Research Grants EY039366, EY019896, and EY024618 from the National Eye Institute (NEI), by Research Grants AI150091, AI143348, and AI147499 from the National Institutes of Allergy and Infectious Diseases (NIAID) to LBM, and by the Research to Prevent Blindness (RPB) grant. This work is dedicated to the memory of the late Professor Steven L. Wechsler “Steve” (1948-2016), whose numerous pioneering works on herpes infection and immunity laid the foundation for this line of research. We are thankful to Dale Long from the NIH Tetramer facility for providing HSV-specific tetramers used in this study.

---

**Supplementary Table1:** Immunodominant T cell epitope peptide positions identified from asymptomatic HSV-seropositive individuals and amino acid sequences from HSV-1 glycoproteins D and B (gD and gB), viral tegument proteins (VP11/12 and VP13/14), and the DNA replication-binding helicase (UL9) proteins.

**Supplementary Figure 1: Example of a gating strategy for analysis of T cell phenotypes in HSV-1-infected mice treated with prime/pull/keep T cell immunotherapy.** (**A**) Representative dot plots show the hierarchical flow cytometry gating strategy. Lymphocytes were first identified based on FSC-A vs SSC-A parameters. Doublets were excluded by gating on FSC-H vs FSC-A to select single cells. Lymphocytes were then gated on CD4 and CD8 expression, and CD8+ T cells were further analyzed for antigen specificity using tetramers corresponding to HSV-1 epitopes: gD_53-61_, gB_183-191_, VP11/12_220-228_, VP13/14_504-512_, and UL9_196-204._ (**B**) Representative flow cytometry plots showing the frequencies of tetramer-positive CD8⁺ T cells and the expression of T_RM_, T_CM_, T_EM_, functional, and exhaustion markers in HSV-1 mice treated with PPK immunotherapy compared with untreated controls. Marker gating within tetramer-positive CD8+ T cells was performed using fluorescence minus one (FMO) controls. PPK **represents** the prime/pull/keep immunotherapy group, and NI represents the non-immunized control group with no treatment or antigen exposure.

**Supplementary Figure 2: Prime/pull/keep T cell immunotherapy does not enhance HSV-1-specific CD8^+^ T cell responses in the spleen.** Frequencies (%) of tetramer-positive CD8^+^ T cells specific for multiple HSV-1 epitopes: gD_53-61_, gB_183-191_, VP11/12_220-228_, VP13/14_504-512_, and UL9_196-204_ were analyzed in the spleen of PPK immunotherapy-treated mice and untreated mice. Experimental groups (P, PP, and PPK) and vaccination timelines are defined in Figure 1. Data are presented as mean ± SEM. Statistical significance was determined using two-way ANOVA followed by appropriate multiple comparisons tests (*∗P<0.05, ∗∗P<0.01, ∗∗∗P<0.001, ∗∗∗∗P<0.0001*; ns, not significant).

**Supplementary Figure 3: Prime/pull/keep T cell immunotherapy does not promote tissue-resident memory (T_RM_) responses in the spleen.** Frequencies (%) of tissue-resident memory (T_RM_) markers were analyzed on gated CD8^+^ T cells isolated from spleen samples of PPK immunotherapy-treated mice and untreated mice. Expression of CD49a, CD69, and CD103 was assessed within gB_183-191_ tetramer-positive CD8^+^ T cells. Experimental groups (P, PP, and PPK) and vaccination timelines are defined in Figure 1. Data represent the mean ± SEM. Statistical significance between the experimental groups was determine using one-way ANOVA followed by appropriate multiple comparisons tests (*∗P<0.05, ∗∗P<0.01, ∗∗∗P<0.001, ∗∗∗∗P<0.0001*; ns, not significant).

## Notes

<u>Support and Conflict of Interest</u>: Studies of this report were supported by Public Health Service Research Grants EY039366, EY019896, and EY024618 from the National Eye Institute (NEI), by Research Grants AI150091, AI143348, and AI147499 from the National Institute of Allergy and Infectious Diseases (NIAID) to LBM, and by the Research to Prevent Blindness (RPB) grant. LBM has an equity interest in TechImmune, Inc., a company that may benefit from the research results and serves on the company’s Scientific Advisory Board. LBM’s relationship with TechImmune, Inc., has been reviewed and approved by the University of California, Irvine, in accordance with its conflict-of-interest policies.

### Competing Interest Statement

The authors have declared no competing interest.

