## Supplementary Table 1 for "A Tissue-Targeted Immunotherapy Protects Against Recurrent Ocular Herpes Infection and Disease, and Alleviates the Associated Cognitive, Behavioral, and Social Deficits in a ‘Humanized’ HLA Transgenic Mouse Model of UVB-Induced Reactivation"

| $CD8^+$ T cell epitopes | Sequences |
|---|---|
| $gD_{53-61}$ | SLPITVYYA |
| $gD_{278-296}$ | ALLEDPVGT |
| $gB_{183-191}$ | GIFEDRAPV |
| $gB_{342-350}$ | NLLTTPKFT |
| $gB_{561-569}$ | RMLGDVMAV |
| $VP11/12_{220-228}$ | RLNELLAYV |
| $VP11/12_{702-710}$ | ALSALLTKL |
| $VP13/14_{504-512}$ | ALHTALATV |
| $VP13/14_{544-552}$ | RLLGFADTV |
| $UL9_{196-204}$ | ALMLRLLRI |
| **$CD4^+$ T cell epitopes** | **Sequences** |
| $gB_{166-180}$ | KDVTVSQVWFGHRYS |
| $VP11/12_{129-143}$ | TQYWKYLQTVVPSGL |
| $gD_{49-82}$ | QPPSLPITVYYAVLERAC<br>RSVLLNAPSEAPQIVR |
