## Supplementary Figure 1 and 2 for "A Tissue-Targeted Immunotherapy Protects Against Recurrent Ocular Herpes Infection and Disease, and Alleviates the Associated Cognitive, Behavioral, and Social Deficits in a ‘Humanized’ HLA Transgenic Mouse Model of UVB-Induced Reactivation"

### Slide 1
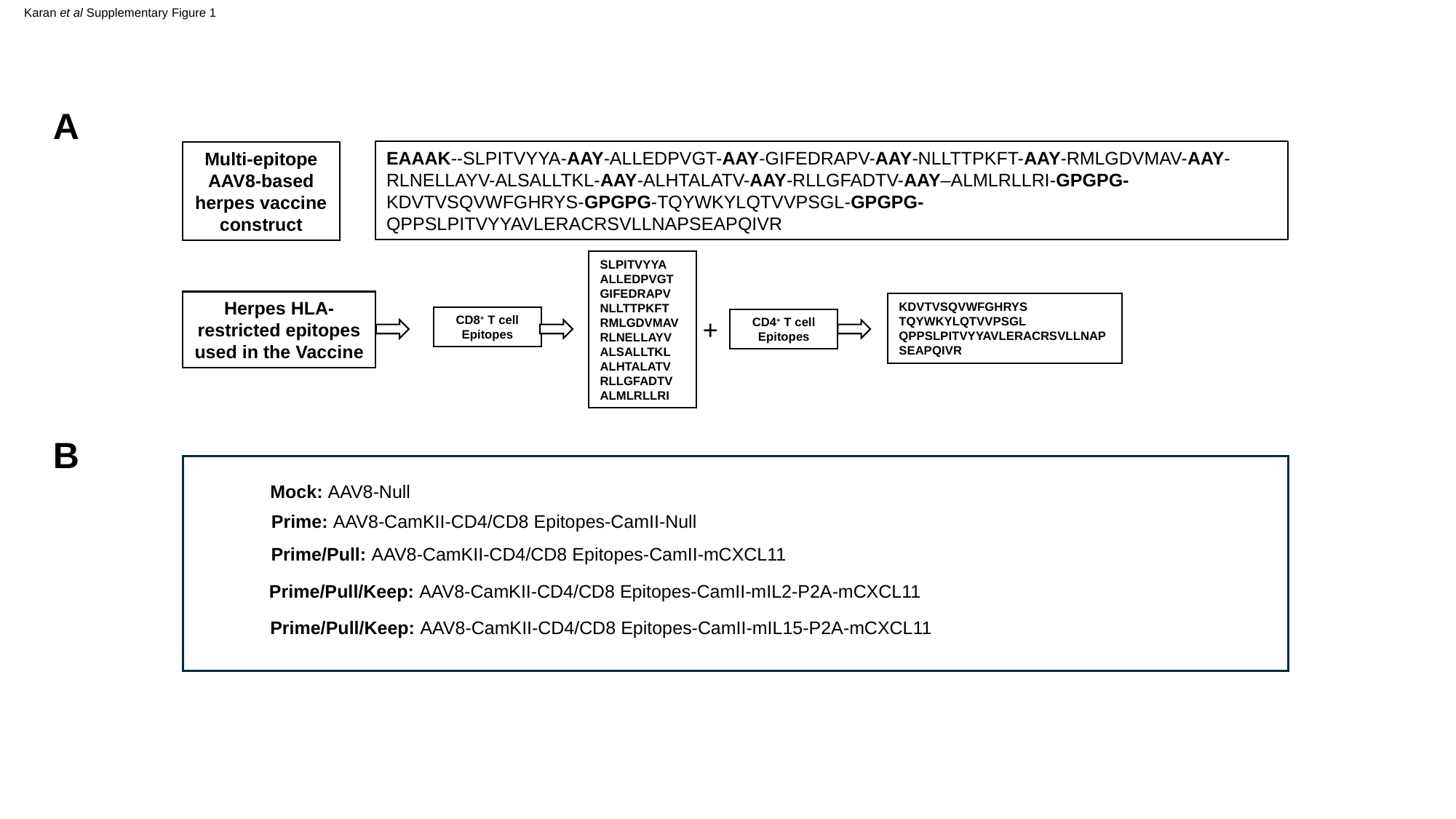

Karan et al Supplementary Figure 1
A
Multi-epitope AAV8-based herpes vaccine construct
EAAAK--SLPITVYYA-AAY-ALLEDPVGT-AAY-GIFEDRAPV-AAY-NLLTTPKFT-AAY-RMLGDVMAV-AAY-RLNELLAYV-ALSALLTKL-AAY-ALHTALATV-AAY-RLLGFADTV-AAY–ALMLRLLRI-GPGPG-KDVTVSQVWFGHRYS-GPGPG-TQYWKYLQTVVPSGL-GPGPG-QPPSLPITVYYAVLERACRSVLLNAPSEAPQIVR
SLPITVYYA
ALLEDPVGT
GIFEDRAPV
NLLTTPKFT
RMLGDVMAV
RLNELLAYV
ALSALLTKL
ALHTALATV
RLLGFADTV
ALMLRLLRI
Herpes HLA-restricted epitopes used in the Vaccine
KDVTVSQVWFGHRYS
TQYWKYLQTVVPSGL
QPPSLPITVYYAVLERACRSVLLNAPSEAPQIVR
CD8^+^ T cell Epitopes
+
CD4^+^ T cell Epitopes
B
Mock: AAV8-Null
Prime: AAV8-CamKII-CD4/CD8 Epitopes-CamII-Null
Prime/Pull: AAV8-CamKII-CD4/CD8 Epitopes-CamII-mCXCL11
Prime/Pull/Keep: AAV8-CamKII-CD4/CD8 Epitopes-CamII-mIL2-P2A-mCXCL11
Prime/Pull/Keep: AAV8-CamKII-CD4/CD8 Epitopes-CamII-mIL15-P2A-mCXCL11

### Slide 2
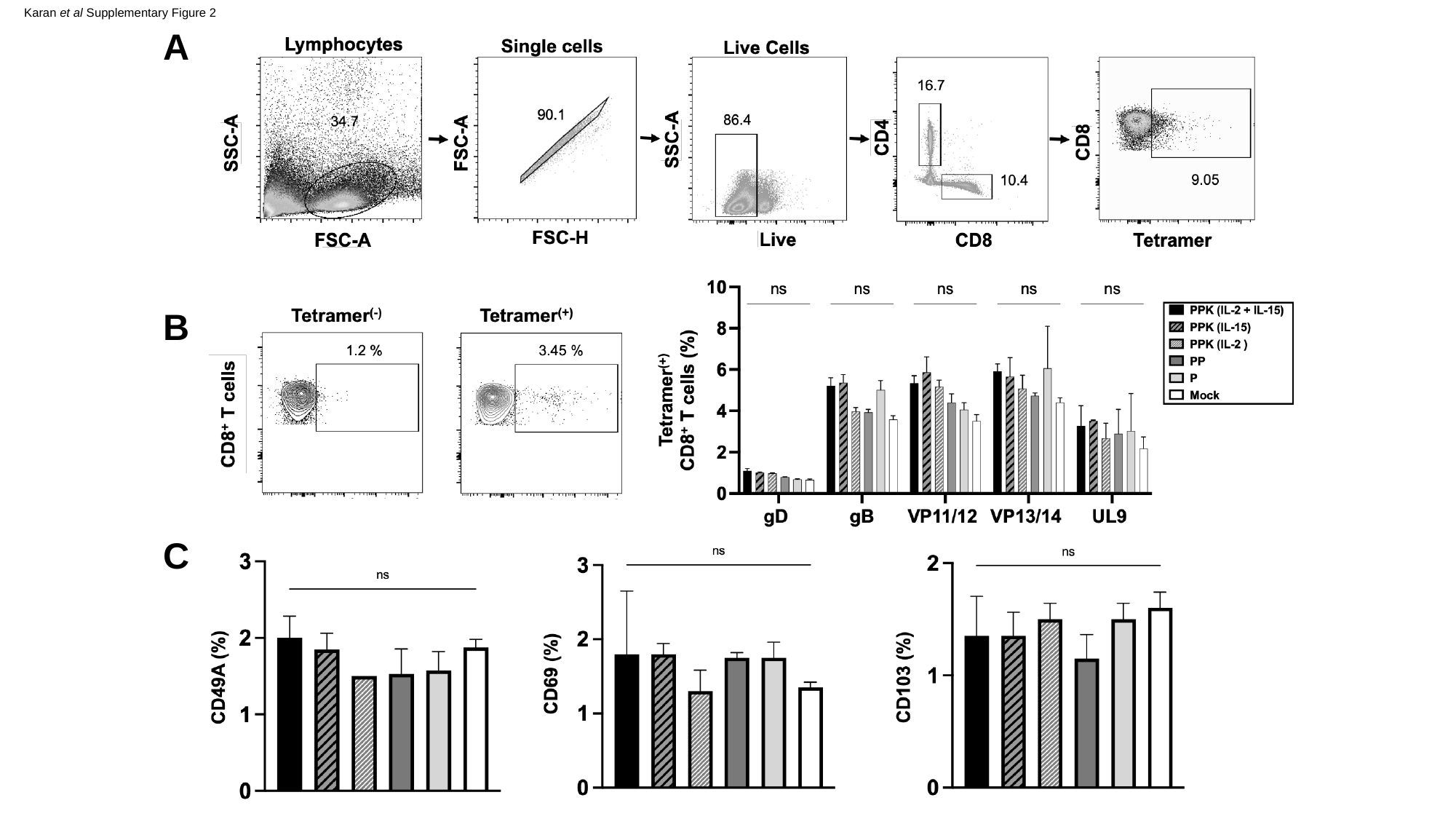

Karan et al Supplementary Figure 2
A
B
C
